# Bioelectricity Generation from Acidogenic Palm Oil Mill Effluents using Microbial Fuel Cells

**DOI:** 10.64898/2026.03.04.709460

**Authors:** Jemilatu Omuwa Audu, Hui Jing Ng, Zaharah Ibrahim, Norahim Ibrahim, Wan Rosmiza Zana Wan Dagang, Mohd Hafiz Dzarfan Othman, Mohd Firdaus Abdul-Wahab

## Abstract

Microbial fuel cell offers a promising approach to improve wastewater quality and generate bioenergy from dark fermented effluents. In this study, the use of dark-fermented palm oil mill effluent as an electron donor for bioelectricity generation was investigated using a double-chambered microbial fuel cell (MFC). The MFCs were operated at room temperature (29 ± 2°C), anode electrolytes adjusted to pH 7, and a chemical catholyte as the oxidizing agent. The maximum power ± 8.07 mW/m^2^ and 155.16 ± 12.88 mA/m^2^, respectively, were generated from the MFCs inoculated with sludge, which was 5.9 times higher than control without inoculum. Microbial community analysis revealed the enrichment of fermentative and electrogenic representative taxa from the phyla *Bacillota*, *Bacteroidota* and *Pseudomonadota* on the anode electrodes. Optimizations of the running conditions were carried out, suggesting the optimum parameters of 0.5 kΩ external resistance, anolyte initial pH 9, and 75% DFPOME substrate concentration. Operation under the optimized conditions increased current production, wastewater treatment, and Coulombic efficiency compared to the non-optimized conditions. Multiple configurations were also evaluated, showing higher cumulative voltage, power, and current densities with the stacked MFC connections, compared to single MFC units. Parallel circuit connection produced higher power and current density than serial connection. This study demonstrated the feasibility of MFC as a promising downstream treatment for biohydrogen production processes, towards higher treatment efficiency and resource recovery.

## 1 Introduction

As industrialization and population grow, energy demand is expected to increase. Current energy sources are based on fossil fuels, and their use pollutes the environment with various contaminants and greenhouse gas emissions. This has necessitated the search for renewable and sustainable alternative energy sources to meet the growing energy demand. Among the promising renewable energy sources, hydrogen is one of the cleanest energy forms, with a calorific value of 122 kJ/g, which is about 2.75 times higher than that of hydrocarbons. In addition, production methods via the process of dark fermentation using various biomass products are currently considered to be an energy-saving approach, with higher production rates and operating procedures more suitable for practical applications [1].

Palm oil mill effluent (POME) is the largest waste generated in some Southeast Asian countries–Malaysia, Thailand, and Indonesia–with humid climates suitable for year-round oil palm production [2, 3]. POME is also one of the most polluting wastewaters with a high organic load, but its high biodegradable content makes it a potential feedstock for biohydrogen production. However, despite progress in improving hydrogen yield by optimizing operating conditions [4, 5], hydrogen-producing community enrichment, and reactor configurations [6], the process still struggles with low COD removal efficiency due to inefficient substrate conversion. The reported COD removal from hydrogen production process using POME as substrate is generally less than 50%. We previously reported COD removal of 35% from a batch mode operation with enriched and acclimated mixed culture inoculum [7]. COD removal of 47.44% was achieved with a pure isolate *Bacillus anthracis* PUNAJAN 1 [8]. Another studies reported COD removal of 48.7% and 35%, with an integrated up-flow anaerobic sludge fixed-film reactor (UASFF) [9, 10], while total and soluble COD removal of 44% and 37%, respectively was achived using ozonated POME [4]. The low substrate conversion is related to the fact that, in addition to the hydrogen produced, a significant portion of the substrates are converted into soluble metabolites that acidify the process and hinder further hydrogen yield [11, 12]. The resulting acidogenic effluent still contains a significant amount of organics, which is typically still above the limits for discharge to water bodies. Therefore, post-hydrogen production processes are needed to improve the quality of the final effluent, and simultaneous energy recovery.

Among the downstream hydrogen processes, anaerobic digestion is the most commonly reported process for secondary treatment of acid-forming effluents from the POME fermentation, with overall total COD removal of over 80% reported for combined processes [13–15]. Increased hydrogen yield with a secondary hydrogen production process has also been studied, for example using a secondary anaerobic sequencing batch reactor operated under thermophilic conditions to improve the total biohydrogen yield, generating from 8.54 to 10.34 mmol H_2_/L/h with a total sugar consumption of 91% [2]. Up to 93% COD removal was achieved with 50% diluted acidogenic effluents via photo-fermentation [8]. Activated sludge process has also been used as a secondary treatment in the use of fermented POME to produce polyhydroxyalkanoates (bioplastics) [16].

In addition to these, microbial fuel cell (MFC) offers a promising approach with the ability to recover energy in the form of bioelectricity, and improve wastewater quality. The increasing interest in the application of MFC in wastewater treatment over the past decade is due to the outstanding benefits of directly converting the chemical energy stored in the substrates into electrical energy without the need for purification or conversion [17]. MFC technologies also offer faster process kinetics, less sludge generation and low energy consumption [11, 18] and insensitivity to a wide range of operating parameters as compared to the conventional anaerobic and aerobic treatment processes [19].

A typical MFC systems consist of an anode and cathode chamber, in which organic-based wastes are oxidized to electrons, protons, and carbon dioxide in the anode chamber. The electrons and protons travel through the circuit and membrane, respectively, to the cathode, generating an electric current [11]. In the cathode chamber, the electrons and protons are reduced to water by oxygen. The preferred substrates for electrogenic microorganisms are organic acids (acetate, lactate, etc.), which are abundant in the acidogenic effluent, making it a suitable substrate for MFC operation [20], the presence of which usually inhibits the traditional fermentation process. Most of the studies conducted so far using POME as an electron donor were performed with dilute POME concentrations with COD value of less than 5,000 mg/L and mostly focused on the performance of a defined electrogenic inoculum, or the comparison of different inoculum types [21–24]. Due to the complex organics in POME, which are generally not easily degraded by most electrogenic bacteria, the POME substrates are pre-hydrolyzed, diluted to appropriate COD concentrations or supplemented with synthetic medium to enhance its bioavailability to bioelectricity [25–28].

The power output of *P. aeruginosa* ZH1 in pure culture and anaerobic POME sludge was compared before, using sterilized POME final effluent with a COD concentration of 2,680 mg/L [24]. A 5.3-fold increase in power density with the pure culture *P. aeruginosa* ZH1 was achieved versus sludge inoculum, but COD removal with the pure culture was only 3% compared to 13% with sludge inoculum. Another study reported that dry sludge produced the maximum power outcome of 3.3 W/m^3^ and 71% COD removal using 1,000 mg/L of COD influent [21]. In an MFC integrated with an adsorption system, the COD concentration of 884 mg/L was utilized and generated a power density of 74 mW/m^2^ and a high COD removal of 93% [29]. The use of sterilized, diluted, or buffer supplemented POME as MFC substrates is associated with high operating costs for large-scale and real-world applications. In this sense, pre-fermentation prior to the electrogenesis steps could make the POME more suitable for MFC systems. However, the use of dark-fermented POME effluent as substrate in MFC systems has not been fully investigated. This study investigated the feasibility of using fermented POME effluent obtained from an ASBR operated under mesophilic conditions [7]. To achieve this, an electrogenic biofilm was first developed in a double-chamber MFC system using static external resistance method in a bufferless system. After stable operation was achieved, the developed system was optimized in terms of the external resistance, anolyte initial pH and substrate concentration. Finally, the performance output was evaluated under serial and parallel configurations.

## 2 Materials and methods

### 2.1 Sample and inoculum preparation

The acidogenic effluents used for the MFC operations were obtained from an ASBR operated for hydrogen production under mesophilic conditions [7] with an HRT of 12 h and OLR of 58.7 g COD/L/d. The characteristics of the acidogenic effluent used are listed in Table 1. POME anaerobic digested sludge was used as inoculum and heat-treated at 90°C for 30 min to inhibit the action of methanogens.

**Table 1.** Characteristics of the dark fermented POME (DFPOME) effluents used in this study.

| Parameters | Results |
| --- | --- |
| pH | 4.35 ± 0.56 |
| Temperature | 35 ± 1.25 |
| Soluble chemical oxygen demand (COD) (mg/L) | 17,350 ± 876.16 |
| Biological oxygen demand (BOD) (mg/L) | 10,510 ± 555.02 |
| Total carbohydrate (mg/L) | 1,950 ± 86.15 |
| Ammoniacal nitrogen (NH<sub>3</sub>-N) (mg/L) | 128 ± 7.50 |
| Total lignin (mg/L) | 155 ± 16.17 |
| Total phenolic content (mg/g of dry mass) | 248 ± 12.89 |
| Color (ADMI) | 3,900 ± 105.45 |
| Acetic acid | 2,100 ± 137.31 |
| Butyric acid | 5,500 ± 175.59 |
| Propionic acid | 550 ± 37.31 |
| Ethanol | 315 ± 18.53 |
| Conductivity (mS/cm) | 4.58 ± 0.78 |

### 2.2 Microbial fuel cell (MFC) design and operation

MFC setups were constructed with two chambers as described before [24] with slight changes in the geometry (Fig. 1). The experimental setups were fabricated into cubic shapes using Plexiglass sheets with dimensions (5 cm × 5 cm × 5 cm) for the anode and cathode chambers. The compartments were separated by Nafion 117 proton exchange membrane (PEM) placed between two rubber sealing gaskets. Three openings of 1, 1, and 0.2 cm diameters were made in the upper part of the two chambers and used as outlets for feed, sampling, and wire ports. Carbon felts with a surface area of 26.3 cm^2^ were used as the electrodes, which were held vertically upright using 1 mm thick copper wires (connecting wires), at 1.5 cm between the electrodes, and connected to an external circuit (1 kΩ external resistance). Before assembling the MFC setups, the PEM was pretreated to increase its conductivity and remove impurities by heating it sequentially in 3%(v/v) H_2_O_2_, 0.5 M H_2_SO_4_, and distilled water at 80℃ for 1 h with each [30]. The carbon felts were soaked in sterile distilled water for 24 h [31] and the remaining components (rubber gasket, butyl rubber, tubing) were autoclaved at 121℃ and 15 kPa for 20 min. the MFC components were assembled aseptically in the laminar flow.

**Fig. 1.**
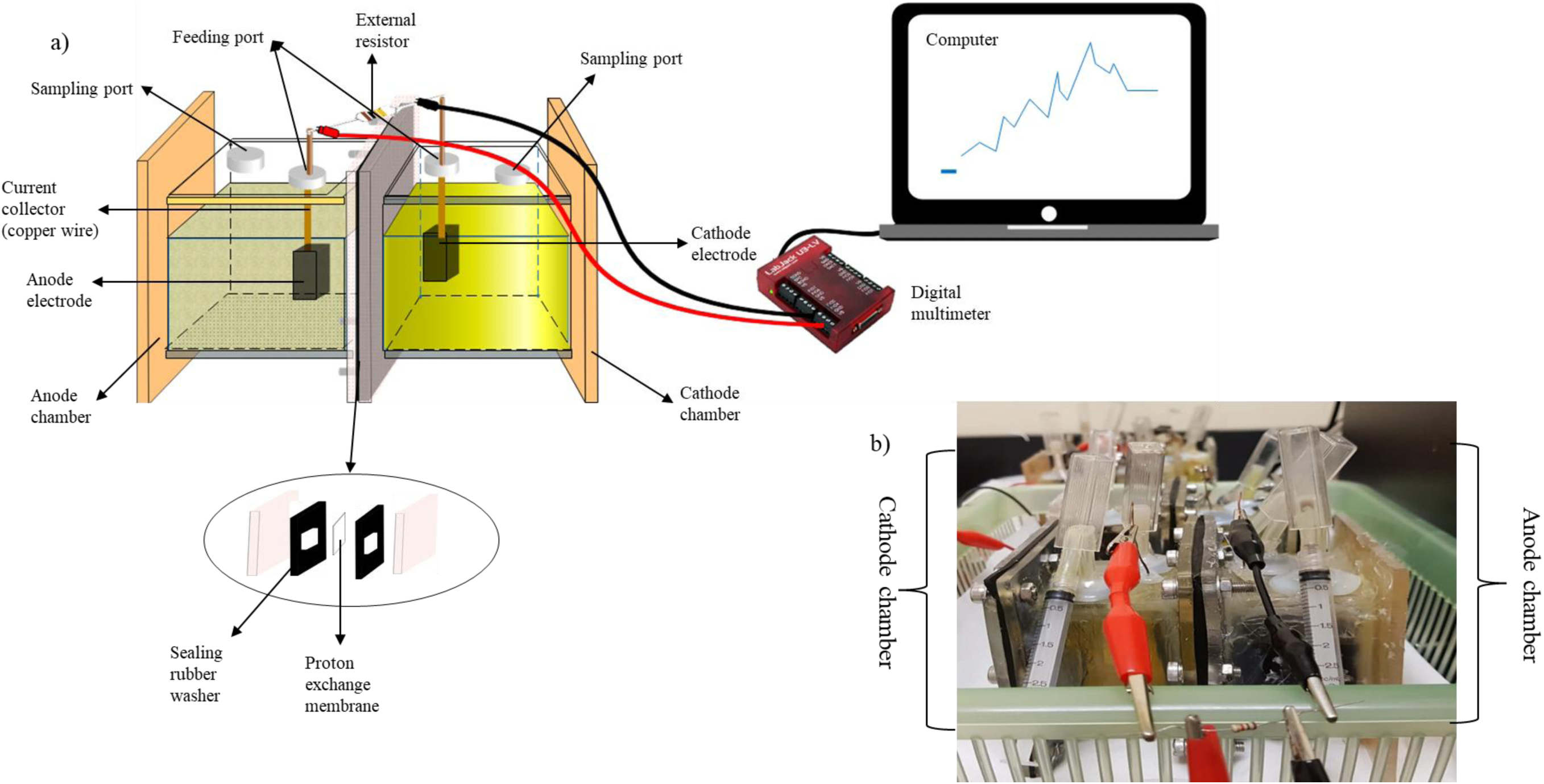
The double-chambered MFC used in this study. (a) Schematic diagram and (b) actual setup.

The MFC systems were operated in semi-continuous mode with a working volume of 100 mL per chamber. The fermented POME (DFPOME) served as the anode electrolyte while 0.1 M potassium ferricyanide(III) (K_3_[Fe(CN)_6_) solution was the cathode electrolyte [21]. The DFPOME was first centrifuged at 4000 rpm for 15 min and the pH was adjusted to 7.0 before inoculating into the anode compartment with 10%(v/v) of sludge inoculum. All MFC setups were operated at room temperature with an external resistance of 1 kΩ to develop and acclimate the electrogenic microbial community. The anode and cathode electrolytes were replaced with fresh solutions when the voltage output dropped below 20 mV, and the catholyte was reduced (yellow to green color) [32]. Two MFC control groups were also established (i) without inoculum and sterilized DFPOME and (ii) DFPOME without inoculum and operated in parallel to confirm that the generated voltage was from the microbially catalyzed and not from purely chemical reactions. The bioanodes were considered developed and stable after three consecutive cycles when relatively stable power output and organic removal efficiency were observed [33].

### 2.3 Power production

MFC performance was evaluated by measuring the potential difference (*V*) across the external resistor (1 kΩ) every 15 min, using a multichannel digital multimeter and a data logger (U3-LV, Labjack). The current (*I*) across the external resistor was calculated using Ohm’s law, and the power (*P*) was calculated from the current and voltage. The power density (*PD*) and current density (*CD*) were determined by normalizing the power and current output to the projected surface area (2.36 × 10^-3^ m^2^) of the anode electrode using Equations 2.1-2.4 [34]. Coulombic efficiency (CE) was calculated as the percentage of total electric current produced over time and divided by the theoretical coulombs that can be produced from the substrates in terms of COD [21] (Equation 2.5).

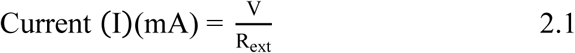

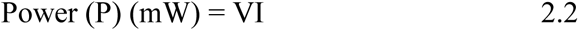

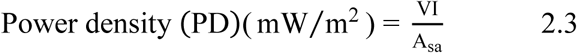

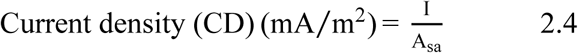

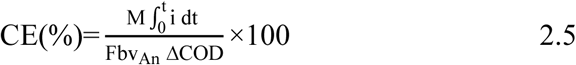

Where V and I are the voltage and current measured across the external resistor (*R_ext_*), and *A_sa_* is the surface area of the anode electrode. M (g O_2_/mol O_2_) is the molecular mass of oxygen, F is the Faraday’s constant (96,485.33 C/mol eˉ), 𝑏 is the number of electrons exchanged per mole of oxygen (4 electrons), 𝑣_𝐴𝑛_ is the volume of liquid in the anode chamber, and Δ𝐶𝑂𝐷 is the change in COD over time.

### 2.4 Electrochemical analysis

The polarization curve measurement was performed at the peak of the cell potential when the maximum voltage was observed and remained relatively stable. The external resistor was disconnected from the MFC and not moved for 2 h before the open circuit voltage (OCV) was measured. Then, the external resistors were varied in descending order (25 kΩ to 51 Ω). Each external resistor was held for 20 min and the voltage across the resistor was recorded [35]. The current, power density, and current density were calculated and used to plot the polarization and power density curves. The internal resistance was determined from the ohmic region of the polarization curve using the polarization slope method. Cyclic voltammetry analysis (CV) was performed under non-turnover conditions at the end of the operated cycles when the oxidizable substrates were presumably depleted [36]. This ensures that the voltammetric behavior is strongly dependent on the applied scan rate (Fricke et al., 2008). The CV measurement was performed within a potential range of -1.0 to 1.0 V using a scan rate of 1 mV/s against a silver/silver chloride (Ag/AgCl) reference electrode.

### 2.5 Effluent analysis

The COD concentration of the influent and effluent was measured using HACH UV-vis spectrophotometer (DR 6000), and the removal efficiency was calculated using Equation 2.6. The composition of volatile fatty acids (VFAs) was determined using gas chromatography equipped with a flame ionization detector (GC-FID) (450-GC, Varian Inc.) with a Restek Stabilwax^®^-DA column. The fermentation effluent was centrifuged at 4000 rpm for 15 min, the supernatant was acidified using 5%(v/v) phosphoric acid, and filtered into 2 mL amber screw cap GC vials using a 0.45 µm filter syringe. The detector, injector, and oven temperatures were set at 250°C, 150°C, and 220°C, respectively. The carrier gas used was hydrogen at a 1 mL/min flow rate. COD removal efficiency was calculated according to Equation 2.6,

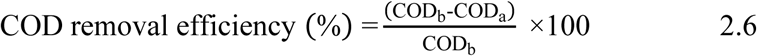

where CODb and CODa are the COD values before and after fermentation, respectively.

### 2.6 Anode biofilm characterization and microbial community analysis

The biofilm formed on the surfaces of the anode electrode was examined using a field emission scanning electron microscope (FESEM) (SU8020, Hitachi). The biofilm was prepared as previously described. The carbon felt electrodes were removed from the MFC setups at the end of the experiments and a piece of the electrode was aseptically excised using a sterile blade. The electrode pieces were rinsed three times with 0.1 M phosphate buffer (pH 7.2) and fixed with a fixative solution containing 2%(v/v) glutaraldehyde and 1%(v/v) formaldehyde solution for 24 h. The excised electrode was then rinsed three times with phosphate buffer and dehydrated in ethanol solutions of increasing concentration (10%, 25%, 50%, 75%, 90%, and absolute ethanol (100%(v/v)) for 30 min for each concentration with regular gentle shaking [37]. The dehydrated electrode pieces were then dried at room temperature and sputter-coated with platinum under vacuum before visualization under FESEM.

Genomic DNA from the sludge inoculum, anode biofilm and planktonic cells was extracted using DNesy^®^ PowerSoil^®^ DNA Extraction Kit (Qiagen) according to the manufacturer’s instructions. PCR amplification, library construction, and amplicon sequencing on an Illumina MiSeq platform were performed by Apical Scientific Sdn. Bhd. (Selangor, Malaysia). The quality of the genomic DNA was estimated by spectrophotometric (Nanodrop) and fluorescence (PicoGreen) methods and visualized using 1.0% agarose gel electrophoresis. The oligonucleotide primers used were illuminaV3/V4F(5’-TCGTCGGCAGCGTCAGATGTGTATAAGAGACAGCCTACGGGNGGCWGCAG-3’) and illuminaV3/V4R (3’-GTCTCGTGGGCTCGGAGATGTGTATAAGAGACAGGA CTACHVGGGTATCTAATCC-5’). The PCR products were purified, quality and quantity measured, and used for library preparation on the MiSeq platform in 2 × 301PE format. The generated sequence length and nucleotide ambiguity was screened by removing sequence shorter than 150 bp and longer than 600 bp during the downstream processing. Chimeric sequence was also detected and removed using the UCHIME algorithm [38].

High-quality reads were analyzed using Quantitative Insights into the Microbial Ecology (QIIME) pipeline and clustered into amplicon sequence variants (ASV) with 97% similarity. The ASVs were classified into different classification levels: domain, phylum, class, family, genus, and species using the SILVA ribosomal RNA database. Alpha diversity, beta diversity, species richness, coverage, diversity (Shannon and Simpson index), and rarefaction analysis were performed using the vegan package in R software v3.5. Statistical significance of the taxonomic units between the samples was assessed using a two-sided Fisher’s Exact test, using XLSTAT 2020.3.1 (Addinsoft) plugin on MS Excel v16.16.24. The representative sequence data for the samples were deposited into the NCBI database under the Bio-project ID PRJNA972304.

### 2.7 Investigation of the factors affecting power performance

After obtaining a stable power production curve for at least three cycles, the effects of the operational parameters on MFC performance were investigated. Single parameter optimization was carried out under different initial anolyte pH, substrate concentration, and external resistance. The selected ranges of the individual factors were based on previous studies [19, 21, 22, 39–50].

The effects of different anolyte pH on MFC power performance were conducted by adjusting the pH of the anolyte to pH 5.0, 7.0, 9.0, and 11.0. The pH range was selected to reflect the acidic, neutral, and alkaline conditions. pH adjustment was done using 5 M HCl and 5 M NaOH. The effects of external resistances were investigated to determine the suitable external resistance in optimising the MFC power performance. The selected external resistance values were 0.1 kΩ, 0.5 kΩ 1 kΩ, and 5 kΩ. The external resistance reflected the external resistance below and above the internal resistance of the setups. The DFPOME was diluted to determine the optimal organic loading rate for current production. The substrate concentration in terms of the COD values of the anolyte represents the amount of oxidisable substrates, which serve as electron donors. The DFPOME was varied by diluting the DFPOME with distilled water to 75%(v/v) (12,750 mg/L), 50%(v/v) (8,500 mg/L), and 25%(v/v) (4,220 mg/L) in addition to the undiluted concentration 100%(v/v) (17,300 mg/L). All the optimisation experiments were conducted under batch mode with freshly replaced electrolytes. Finally, the MFCs were operated in a stacked and semi-continuous manner mode, achieved by electrically pairing three single MFCs in serial and parallel arrangements. The voltage reading of the stacked MFCs was monitored across the external resistance using the Labjack multi-channelled data logger.

## 3 Results

### 3.1 Power generation

All MFC systems were operated for nine cycles. The first cycle after inoculation showed no significant increase in power generation and might be related to substrate consumption for cell proliferation (Fig. 2a). Current production started after the first feed change (2nd cycle), and reached a peak power density of 4.38 ± 0.83 mW/m^2^ for MFC-S and 1.25 ± 0.12 mW/m^2^ (MFC-WS) at the 32^nd^ and 74^th^ h, respectively. Thereafter, the power output decreased, indicating the depletion of readily degradable substrates in the anolyte [21]. In subsequent cycles with MFC-S and MFC-WS, a gradual increase in current production was observed after each replacement of the anolyte (feed). This indicates increased biological activity as bacterial cells colonize the anode electrode and enhance the electron transfer process [29]. The MFC-S showed reproducible cycles (<10%) in the last four cycles with an average power density of (60.36 ± 14.66 mW/m^2^). Also, the relatively stable periods increased from 42 h (fourth cycle) to 92 h (ninth cycle), corresponding to an increase in the cycle duration from seven to ten days. The increased stable periods may be an indicator of successful bacterial attachment on the surfaces of the anode electrodes and degradation of the complex organic matter into a variety of easily utilized substrates by the electrogenic microbial community in the system.

**Fig. 2.**
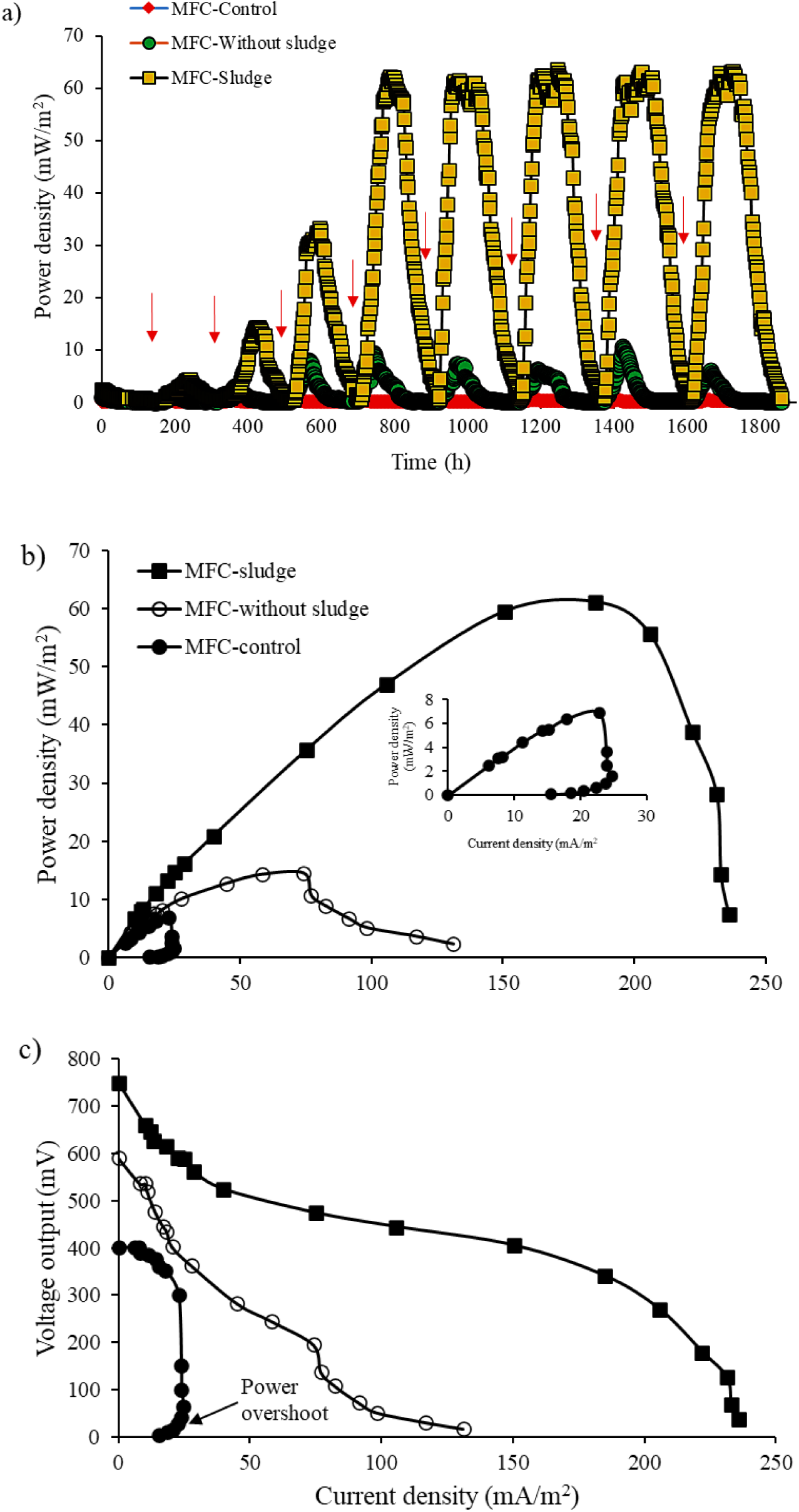
(a) Power density profile of the three MFC setups during the electrogenic community enrichment, (b) power density, and (c) voltage-current curves under different external loads (25 kΩ to 51 Ω). Arrows indicate the feed replacement period

On the other hand, MFC-WS showed fluctuations in power production with a much shorter cycle duration of about five days and a more drastic decline in voltage output. The maximum power density obtained with MFC-WS systems is almost six times (10.65 ± 2.55 mW/m^2^) lower than that obtained with MFC-S. This can be attributed to fewer redox activities at the anode caused by the absence of added inoculum [51]. The current generation of MFC-C remained constant (0.05-0.57 mW/m^2^) throughout the acclimation process, with no significant increase in voltage generation in all the cycles operated.

Polarization measurements conducted at the end of the acclimation period also confirmed the benefit of added inoculum. The generated open circuit voltage (OCV) was 750.04 ± 24.29 for MFC-S, 591.14 ± 36.01 for MFC-WS, and 410 ± 8.01 mV for MFC-C (Fig. 2c). For MFC-S, the voltage readings dropped linearly from 750.04 ± 24.29 to 340 ± 28.87 mV as the current density increased toward 200 mA/m^2^. The current density eventually reached 235 mA/m^2^ with the output voltage decreasing due to the high current demand at low external resistance. The linear region observed in the voltage curve of the MFC-S is probably due to ohmic losses due to the resistance of the fuel cell components and the proton transport process [21]. MFC-WS showed a sharp voltage drop at a current density of less than 150 mA/m^2^ and was likely dominated by activation loss due to insufficient energy required to drive the oxidation-reduction redox reaction at the anode surface [52, 53]. MFC-C showed no significant increase in current density with the changes in external loads due to the absence of biocatalysts. In addition, power overshoot was also observed, noted by the doubling-back effect at the current density (<30 mA/m^2^) with the applications of lower external resistances. This can be attributed to the fact that no or insufficient biofilm developed on the electrode surfaces of MFC-C [54]. From the power density curves (Fig. 2b), the maximum achievable power density is 61.18 ± 6.49 mW/m^2^ for MFC-S, 14.47 ± 3.43 mW/m^2^ (MFC-WS), and 6.87 ± 0.94 mW/m^2^ (MFC-C) at a current density of 184.95 ± 19.92 mA/m^2^ (MFC-S), 74.17 ± 12.03 mA/m^2^ (MFC-WS), and 22.85 ± 2.29 mA/m^2^ (MFC-C). The internal resistance calculated from the ohmic region of the polarization curves showed that MFC-S has the lowest internal resistance of 524 Ω, while MFC-WS and MFC-C have a much higher internal resistance of 1368 Ω and 1520 Ω, respectively.

### 3.2 COD removal and Coulombic efficiencies

The COD removal increased with MFC-S and MFC-WS, as the cycles progressed and was relatively stable over the last four cycles (Fig. 3). The maximum COD removal of 29.22 ± 2.01% for MFC-S and 22.98 ± 2.84% for MFC-WS was observed. COD removal for MFC-C was negligible (0.17 ± 0.51%), probably due to chemical degradation. The CE, which measures the number of electrons converted to electricity from the substrates oxidized, showed that more electron conversion occurred in MFC-S than in MFC-WS, with the maximum CE of 6.89 ± 0.89% in MFC-S and 1.5 ± 0.97% in MFC-WS. Despite the relatively similar COD removal efficiencies of MFC-S and MFC-WS, the reducing equivalents of MFC-WS may have been used to form metabolite, as evidenced in the resulting effluents with a more acidic pH of 4.9 ± 0.25. The drop in pH indicates the accumulation of volatile fatty acids (VFAs), indicating non-electrogenic metabolisms [55]. In contrast, the pH of MFC-S effluent was slightly acidic (6.26 ± 0.19), indicating that a greater portion of the reducing equivalent was used for electrogenesis as confirmed by the higher CE. The lower CE of the operation without sludge indicates the non-electrogenic removal of organic matter by fermentative microorganisms in the DFPOME effluents. Although the CE of MFC-S is lower than the values reported for the operation pure culture [23, 56], it is similar to the values reported with mixed culture [22, 57]. The low CE is a common phenomenon in real wastewater utilization due to the high activities of the other metabolic pathways (biomass formation, metabolite production, and possible alternative electron acceptors (sulfate and nitrate), that divert the electrons available for the anode electrode surfaces [53, 58].

**Fig. 3.**
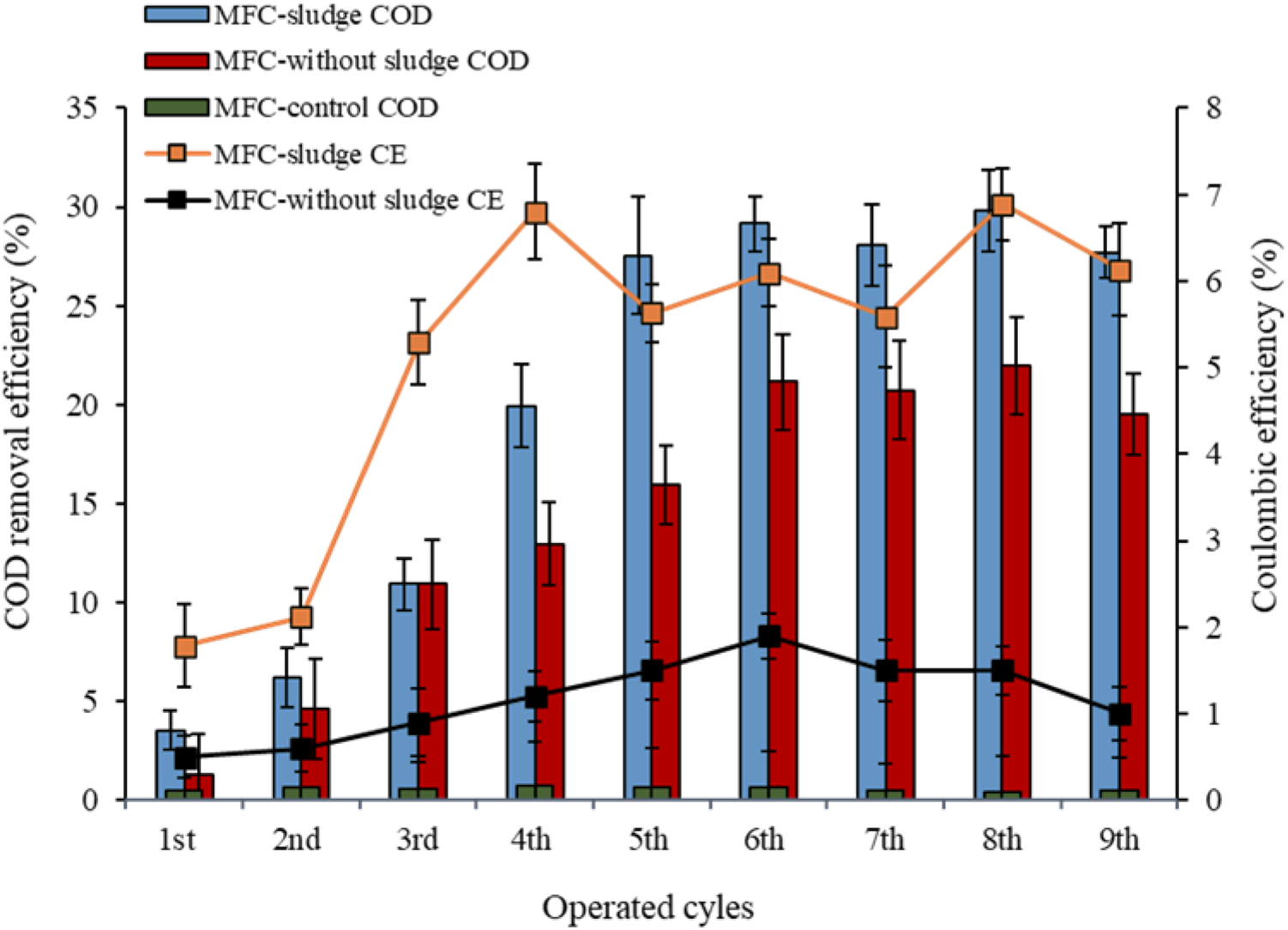
COD removal and Coulombic efficiencies of the three MFC setups during the enrichment of the electrogenic community

### 3.3 Cyclic voltammetry (CV) analysis

The electrochemical reactions at the bacterial or biofilm-electrode interfaces were elucidated using cyclic voltammetry. A catalytic current of 0.023 A was observed for MFC-S and 0.013 A for MFC-WS of the oxidation and reduction peaks, whereas MFC-C exhibited no electrochemical activity due to the absence of redox peaks on the cyclic voltammogram (Fig. 4). The two redox couples were observed for MFC-S, with the most prominent couple at the formal potentials of 0.281 V (denoted as *a_ox_*) and 0.183 V (*a_rd_*), with current peak heights of 0.015 A and 0.008 A, respectively. The redox couple at the indicated potential may be due to the cytochrome complex of the bacterial cell surface reacting with the anode electrode. The less prominent couple at the negative formal potentials of -0.403 V (*c_ox_*) and -0.347 V (*c_rd_*), most likely corresponds to the electron carriers NAD^+^/NADH and NADP^+^/NADPH (Kracke et al., 2015). Although the current peak of the latter redox couple is lower, it could still indicate participation in the electron transport processes. Additional oxidation peaks were observed at the formal potentials of 0.696 V (*b_ox_*) and -0.559 V (*d_ox_*) without the corresponding reduction peaks. The first peak could be attributed to the oxidation of oxygen present in the anode chamber, while the second peak could be due to unidentified metabolic reactions. In contrast to MFC-S, the redox couple observed in the cyclic voltammogram of MFC-WS was at 0.439 V (*e_ox_*) (0.006A) (Fig. 4), with a much lower correspondingly reduced peak (0.001 A) at 0.647 V. An additional reduction peak without the corresponding oxidation peak was observed at - 0.725 V (*f_rd_*). This could be due to the possibility of a reduced inorganic compound from the metabolic reactions or substrates used as the electron donor. The cyclic voltammogram supports the power production curves, and shows that current production was higher with MFC-S than with MFC-WS. The quasi-reversible and irreversible redox reactions shown in the voltammograms are the characteristics of fermentative electrogens [59] and MFC systems operated with complex substrates [60].

**Fig. 4.**
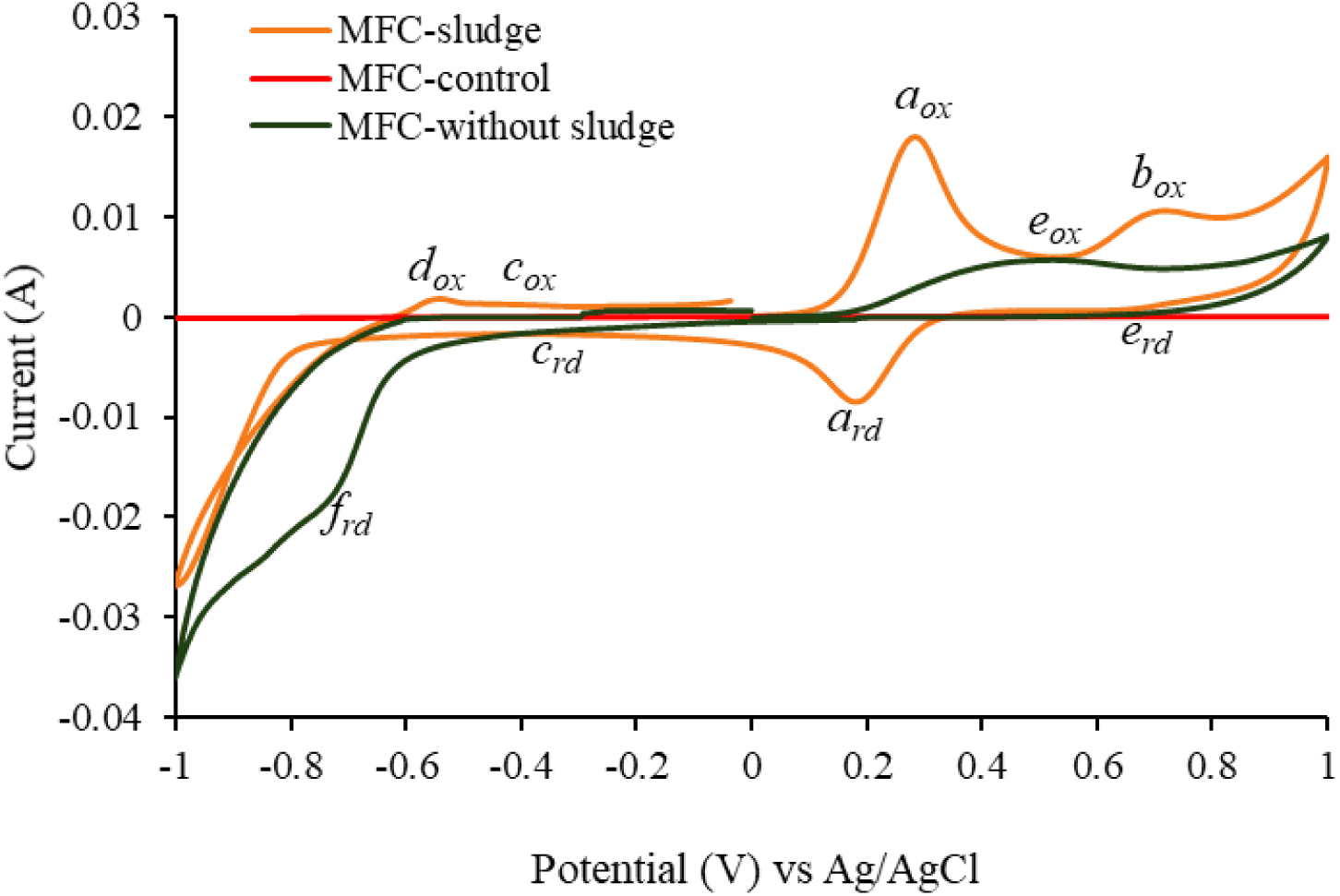
Cyclic voltammograms for the different MFC setups. Oxidative peaks are labeled *aox*, *box*, *cox*, *dox*, *eox*, while the reductive peaks are labeled *ard*, *crd*, *erd,* and *frd*

### 3.4 Analysis of the anodic biofilm

Due to the significant power production and Coulombic efficiency of MFC with added sludge compared to MFC without sludge, the MFC-S systems were considered for further analysis. The biofilm formed was visualized by FESEM. The carbon-felt electrodes were analyzed before and after the acclimation period. Compared to the carbon felt electrode before the MFC operation (Fig. 5a), the FESEM microphotographs of the electrodes with biofilm showed good colonization with biomass consisting of bacterial cells and extracellular polymeric substances (EPS) (Fig. 5b). Clumps of bacterial cells were observed at some locations on the anode surface (Fig. 5c), previously reported to be fermentative bacteria that hydrolyze the complex organics in the POME substrates, providing substrates for the electrogenic bacteria [21]. Rod-shaped bacterial cells with lengths between 0.9-2.1 μm and width of 0.2-0.6 μm dominated the bacterial consortium. At higher magnification (40,000×) web-like structures were visible that could have contributed to the attachment of the bacterial cell to the anode surfaces (Fig. 5e). These structures could also include conductive pili that serve cell aggregation and may be involved in the extracellular electron transfer process [61]. Overall, the FESEM images indicate biofilm formation on the carbon felt electrode.

**Fig. 5.**
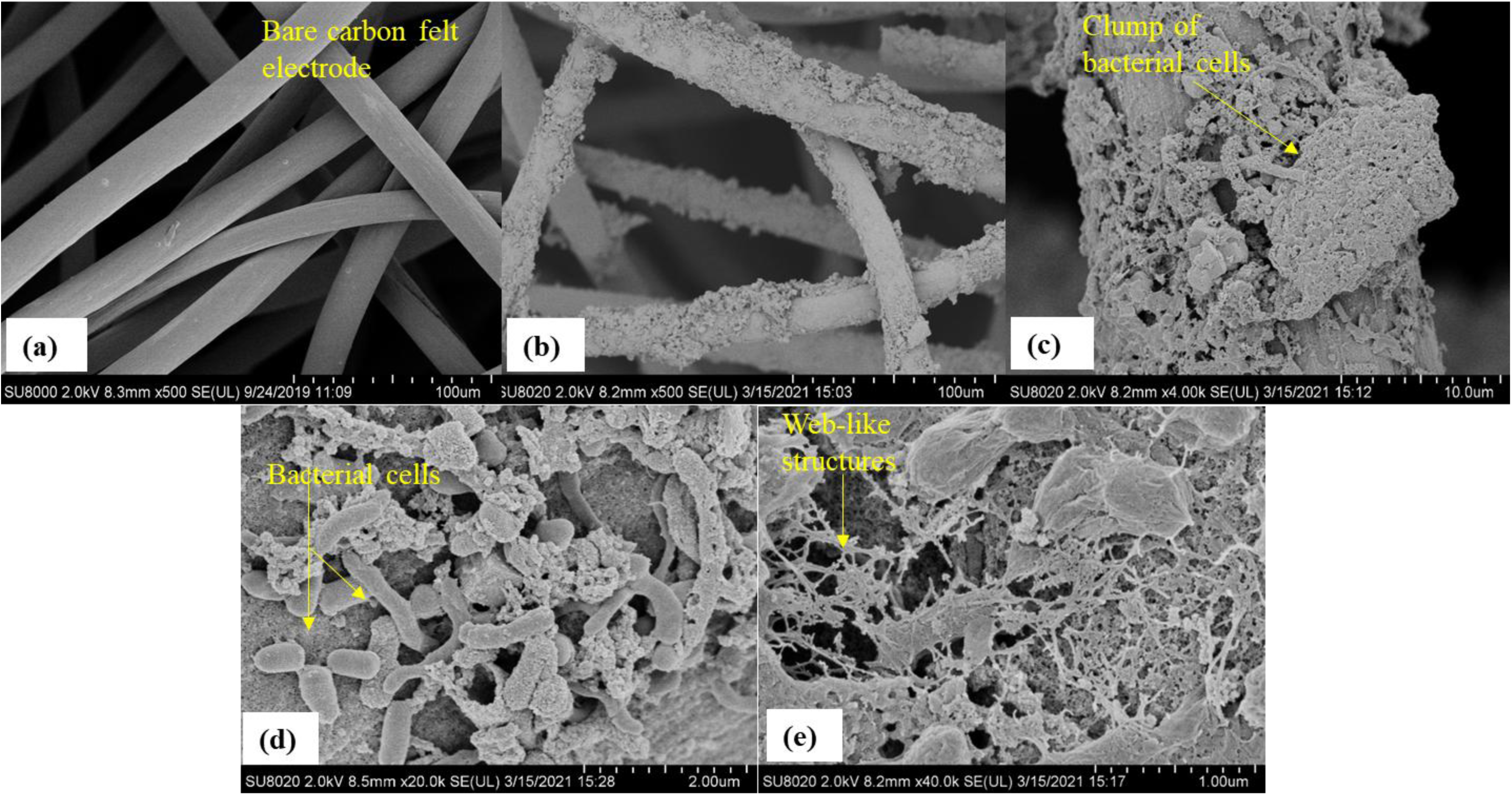
FESEM micrographs of (a) control electrode (bare carbon felt), (b) 500×, (c) 4,000×, (d) 20,000×, and (e) 40,000× magnification of the biofilm formed on the electrode surfaces

### 3.5 Microbial community analysis

The microbial communities in the sludge before the operation, anodic biofilm (AB), and planktonic cells (PC) after the acclimation phase were analyzed using the high throughput 16S rRNA gene sequencing. The ASVs identified with 97% nucleotide similarity revealed observed features of 474.33 ± 51.47 for the sludge sample, 300.33 ± 10.69 (AB), and 222 ± 30.61 (PC). The dominant phyla of the sludge sample were *Bacilliota* (31.92 ± 2.43%), *Bacteriodiota* (26.52 ± 0.89%), *Chloreflexota* (8.14 ± 0.72%), phyla which representing the Archaea domain (*Halobacteriota*, *Methanobacteriota*, *Nanoarchaeota*, *Thermoplasmatota*, and *Thermoproteota*) (8.17 ± 2.86%), *Cloacimonadota* (7.38 ± 0.19%), *Halobacteriota* (4.93 ± 2.42%), and *Spirochaetota* (2.69 ± 0.33%) (Fig. 6). The remaining phyla had a relative abundance of (<2%) of the total sequences. Archaeal phyla were not detected within the anodic biofilm and planktonic cells, indicating the successful suppression of the methanogenic community by the heat pre-treatment method used. A significant difference was also observed in the microbial community of the anodic biofilm and planktonic cells with the enrichment of *Bacillota* phylum to 50 ± 7.09% and 62 ± 11.93%, respectively. *Pseudomonadota*, *Actinomycetota*, and *Desulfobacterota* were also enriched to 9.33 ± 3.51%, 5.67 ± 2.31%, 6.33 ± 0.58% for anodic biofilm and 4.33 ± 3.79%, 6.33 ± 1.53%, and 8.67 ± 3.21%, for planktonic cells. In contrast, *Bacteroidota* reduced to 19 ± 2.31% and 24.67 ± 5.29% for anodic biofilm and planktonic cells, respectively. All other phyla were repressed or reduced to less than 0.1% of the total bacterial sequences (Fisher’s Exact test, P < 0.05) (Fig. 6a).

**Fig. 6.**
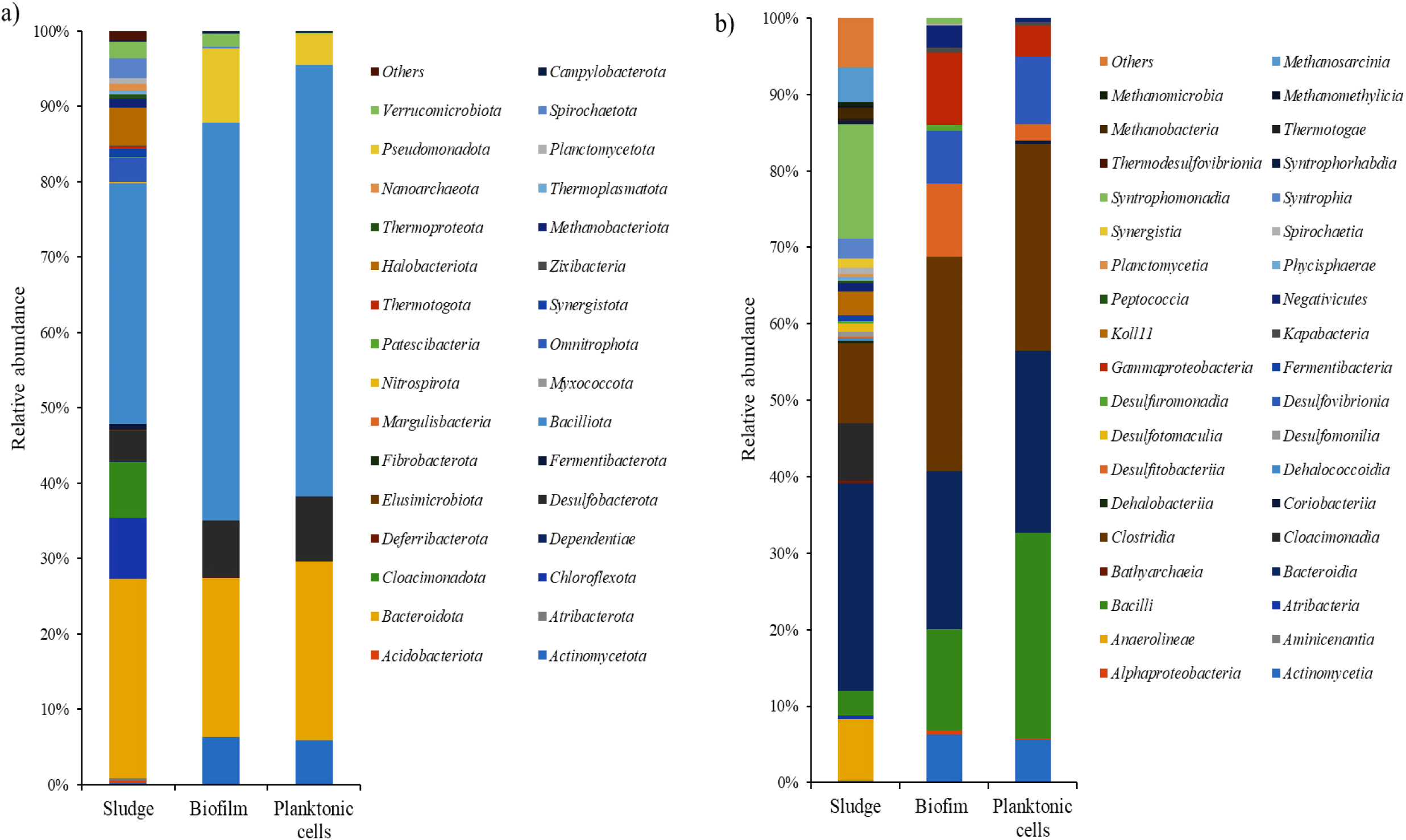
Relative abundance of the microbial community in the sludge, anodic biofilm, and planktonic cells at the phylum (a) and class (b) levels

The class level distribution (Fig. 6b) showed that *Bacteroidia* (26.5 ± 0.88%), *Syntrophomonadia* (14.6 ± 2.29%), *Clostridia* (10.2 ± 0.45%), *Anaerolineae* (7.88 ± 0.75%), and *Cloacimonadia* (7.38 ± 00.19%) were the most abundant in the sludge sample. After MFC operations, *Clostridia* was the most abundant in biofilm and planktonic cells, with an increase in relative abundance to 35.34 ± 3.51% and 31.33 ± 0.57%, respectively. This was followed by *Bacteroidia* (18.67 ± 4.04%), *Bacilli* (11.67 ± 2.08%), *Gammaproteobacteria* (8.33 ± 2.54%), and *Actinobacteria* (5.67 ± 2.31%) in biofilm. Within the planktonic cells, the most abundant classes after *Clostridia* were *Bacilli* (25.67 ± 11.55%), *Bacteroidia* (24.33 ± 0.58%), and *Actinobacteria* (6.33 ± 1.53%). *Gammaproteobacteria* was less than 4%. In addition, *Syntrophomonadia*, *Desulfuromonadia*, *Spirochaetia*, and *Limnochordia* were found only in the biofilm, but in low relative abundance (<1%). *Clostridia* and *Bacteroidia* were observed to be the dominant class in this study, as in several previous studies using complex wastewater [41, 51, 62, 63]. Their abundance can be attributed to the ability of members of this class to hydrolyze organic matter, which is further used by electrogens to generate electricity [41, 51]. *Actinobacteria* are also known to be involved in the degradation of organic pollutants and have been found in an MFC system [64]. Within the *Pseudomonadota* phylum, only the *Gammaproteobacteria* and *Alphaproteobacteria* classes were enriched. In contrast to several other studies, the *Deltaproteobacteria* class, which includes the genera *Geobacter,* was not enriched in this study, likely due to the different enrichment method, electron donor, or the sludge material used and inoculum.

At the genus level, most of the sequences assigned to *Clostridia* belonged to the genera *Clostridium_sensu_stricto_1* of the family *Clostridiaceae*, with a relative abundance of 9% (AB) and 11% (PC). Its presence in both biofilm and anolyte indicates that substrate oxidation occurred in both phases. In addition to their fermentative nature, several *Clostridium* species have been reported to directly participate in current production. *C. butyricum* possessed membrane-bound cytochromes used for direct electron transfer [65]. Other species of the same genus may have similar electron transfer mechanisms [41, 64]. Other reported electrogenic species include *C. beijerinckii* [66], *C. butyricum* [65, 67], *C. acetobutylicum* [68]. *Desulfitobacterium,* of the family *Desulfitobacteriaceae* was found in higher abundance (8 ± 3.21%) in the biofilm than in the planktonic community (2.23 ± 1.33%). Members of the genera (*D. hafniense*) have been reported to participate in electricity generation in an MFC [69] using humic acid as an electron shuttle [70]. Humic acid in POME effluent might have been utilized in this way. Within the *Bacilli* class, the most represented genera were *Bacillus*, *Lactobacillus*, *Limosilactobacillus*, and *Rummeliibacillus*, with a relative abundance of 4.1-8.6% and were more abundant in planktonic cells than in the anodic biofilm. These genera are known hydrolyzers due to their ability to secrete various metabolic enzymes to degrade complex polymers (carbohydrates, protein, lipids, etc.) [71].

Members of the *Bacteroides,* also known as fermenters, are also usually found in MFC systems operated with complex substrates. In this study, a higher relative abundance of *Bacteroides* was found in the planktonic cells (10 ± 4.58%) than in the anode biofilm (3.33 ± 1.53%). Like *Clostridia*, they are widely distributed in organic-based MFC systems due to their hydrolytic and fermentative capabilities. *Bacteroides* are acetate producers, with some members reported to have the ability to transfer electrons by Fe(III) reduction [21, 72]. Their Fe(III) reducing abilities and their presence in the anode biofilm of this study may indicate their involvement in the electron transfer process. In a previous study, *Bacteroides* sp. W7 was isolated from the anodic suspension of a double-chambered MEC system [73]. The co-dominance of *Bacteroides* and *Geobacter* as the dominant genera after three cycles of MFC operation with POME using dry sludge as inoculum has been reported [21]. Another member of the *Bacteroidia* class, *Dysgonomonas*, may have been involved in current production due to its higher relative abundance (4.0 ± 1.0%) in the biofilm than in the planktonic cells (1.0 ± 0.87%). Although the electron transfer capabilities of *Dysgonomonas* sp., are yet to be determined in MFC systems. Some members of the genus have been reported to be associated with increased power output in an MFC system treating a mixture of organic wastes [74, 75].

A higher abundance of *Pseudomonas* (5.33 ± 1.15%) was also found in the anodic biofilm than in the planktonic cells. *Pseudomonas* are Gram-negative facultative anaerobes that produce electron shuttles, such as pyocyanin and phenazines, contributing to MFC performance [76]. The predominance of *Pseudomonas* in the anode biofilm over the planktonic cells suggests their involvement in current production. Interestingly, *Hydrogenophaga,* another genus of interest within the *Pseudomonadota* phylum of the family *Comamonadaceae*, was also detected within higher abundance (4.21 ± 1.05%) in the anodic biofilm than in planktonic cells. *Hydrogenophaga* is a known hydrogen-consuming electrogenic bacteria (Kang et al., 2017), and its presence in the biofilm suggests indirect interspecies electron transfer might have occurred with other electrogens [36]. Such syntrophic cooperation was previously suggested in studies using *Geobacter* [27, 36]. Sulfate-reducing bacteria (*Desulfovibrio*) was observed to be higher in the planktonic cells (9.33 ± 2.08%) than in the anode biofilm (6.0 ± 0.58%). This indicates that the removal of sulfur contaminants occurred mainly in the electrolyte [77]. *Desulfovibrio* has been reported among the microbial community in previous MFC operations, and its presence likely affects the current generation process by using sulfate as the terminal electron acceptor instead of the anode electrode [78]. However, some members (*D. desulfuricans*) have been reported to perform direct extracellular electron transfer to the anode surfaces, contributing to electricity generation [30, 79].

### 3.6 Factors affecting power performance

#### 3.6.1 External resistance

Once relatively stable operations were obtained, the effect of different operating factors (initial anolyte pH, external resistance, and substrate concentrations) were investigated. Three MFCs were used for each factor for the optimization and operated in batch mode. The biofilm-developed electrode was used as the anode electrode, electrolytes (DFPOME and K_3_[Fe(CN)_6_]) were replaced with fresh medium for all conditions and operated in three repeated batch cycles.

Several different external resistances were used (0.1 kΩ, 0.5 kΩ, 1 kΩ, and 5 kΩ), while the anolyte pH and organic load were maintained at pH 7 and 19,300 mg/L (undiluted DFPOME). The box plots show that as the external resistance increases, the voltage exerted by the system also increases (Fig. 7a). MFC-5kΩ showed wider voltage data dispersal, indicating a much wider voltage output variation compared to MFC-0.5kΩ and MFC-1kΩ. The maximum voltage output of the different operated external resistance was as follows: 85.48 ± 28.63 mV (0.1 kΩ), 287.87 ± 39.52 mV (0.5 kΩ), 409.01 ± 28.37 mV (1 kΩ), and 749.07 ± 83.45 mV (5 kΩ). External resistance contributes to the ratio between voltage and current generated. Higher external resistances exhibit higher potential difference, and vice versa [53, 80]. However, the electron flux from the anode to the cathode is hindered by higher external resistance. As observed in this study, the current density generated was inversely proportional to the external resistance applied (Fig. 7b). Current density increases as the applied external resistance decrease, with MFC-0.1kΩ having the highest current density of 342.01 ± 9.28 mA/m^2^. However, the current density generated with 0.1 kΩ was only sustained for 25 h and was followed by a rapid decline resulting in a final current density between 57.98–61.03 mA/m^2^. MFC-0.5kΩ exhibited a relatively longer steady phase current (62 h) before the decline in current output. Despite generating lower current densities, operations with higher external resistance (MFC-1kΩ and MFC-5kΩ) had a much longer plateau of 86 h and 93 h, respectively (Supplementary Figure S1). These results suggest that the metabolic activities of the electrogenic microbial community in the MFC contribute to stable current production irrespective of the external resistance introduced [42].

**Fig. 7.**
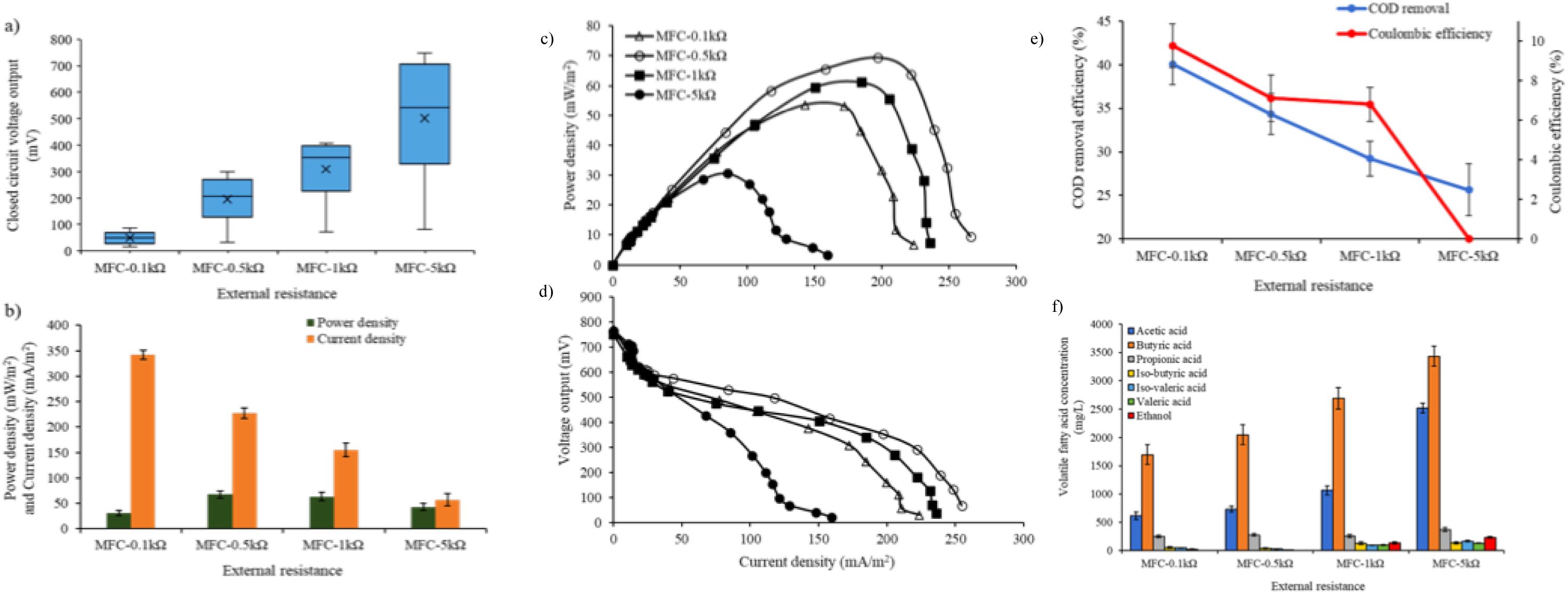
MFC power performance under the different external resistances (0.1, 0.5, 1, and 5 kΩ), (a) closed circuit voltage, (b) maximum power and current density, (c) power density-current curves, (d) voltage-current curves, (e) organic matter (COD) removal and Coulombic efficiencies, and (f) metabolite profile

Operation with low external resistance has been reported to increase the anode potential, which is a measurement of the anode electrode charge, determining the flow of electrons to the circuit [50]. The more pronounced increase in current generation observed with MFC-0.1kΩ and MFC-0.5kΩ can be attributed to the increased rate of electron transfer to the circuit due to the less obstruction of the electron flow with the low external resistance applied [81]. Despite the high potential difference observed with 5 kΩ, the electron flux between electrodes was significantly hindered. This suppresses anode respiration, resulting in the diversion of dominant metabolic pathways to fermentation (acidogenesis) [46]. The results suggest that external resistance of more than 1 kΩ becomes the rate-limiting step in the current generation process and is not favorable for power production [39]. The results observed here align well with the previous studies looking into the effects of external resistance [39, 50]. Comparable power density was obtained with both external resistance of 1 and 0.5 kΩ. Further decrease of the external resistance to 0.1 kΩ resulted in a 2-fold (30.76 ± 4.82 mW/m2) decrease in power density, while the increase to 5 kΩ resulted in a 1.5-fold (42.67 ± 7.07 mW/m^2^) decrease. The power density curves (Fig. 7c) also show that the maximum power attainable with the system was at 0.5 kΩ external resistance (69.39 mW/m^2^). The internal resistances (ohmic loss) calculated from the polarization voltage-current curves (Fig. 7d) are: 643 Ω for MFC-0.1kΩ, MFC-0.5kΩ (560 Ω), MFC-1kΩ (543 Ω), and MFC-5kΩ (1478 Ω). MFC-0.5kΩ and MFC-1kΩ had the lowest internal resistance, while the higher ohmic loss observed with MFC-0.1kΩ and MFC-5kΩ might have contributed to the variation in power output. The ohmic loss of MFC-0.5kΩ and MFC-1kΩ systems in this current study was similar, which could be attributed to the similar power output. However, the power density obtained with MFC-0.5kΩ was slightly higher than that of MFC-1kΩ.

COD removal increased by more than 1-fold when the external resistance was decreased to 0.5 kΩ and 0.1 kΩ, respectively (Fig. 7e). This suggests that the higher current generation obtained with lower external resistances resulted in increased substrate oxidation. When the external resistance was increased to 5 kΩ, a further 1-fold decrease in COD removal was observed, suggesting the decrease of substrate degradation and the possible shift of the bacterial metabolism towards more metabolites (i.e. VFAs) production [46]. Likewise, the Coulombic efficiency increased with the decrease in external resistance, with MFC-0.1kΩ yielding the highest CE of 9.75 ± 1.09% (Fig. 7.3). MFC-0.5kΩ and MFC-1kΩ had comparable (T-test: P > 0.05) CEs of 7.10 ± 1.17% and 6.89 ± 0.86%, respectively. MFC-5kΩ operation drastically decreased CE to 0.02 ± 0.02%, indicating the negative influence of the high external resistance probably due to a change in bacterial metabolisms [50]. The suppression of anode respiration has been reported when the double-chambered MFC was operated with 10 kΩ external resistance [46]. The electron flux from the anode biofilm to the electrode surface was hindered, resulting in increased acetate production (fermentation). The COD removal and Coulombic efficiency obtained in this study are comparable with previous studies, reporting that the increase in external resistance affected the COD removal efficiency and CE output [42, 82]. External resistance was found to influence metabolite production and utilization, with the knowledge that these soluble metabolites include those from the DFPOME substrate used and those produced during the oxidation process in the anode chamber (Fig. 7f). Butyric and acetic acid was the main metabolites detected at all the external resistance used, while the remaining metabolites (propionic acid, isobutyric acid, isovaleric acids, valeric acids and ethanol) were detected in lower concentrations (<250 mg/L). Also, ethanol was only detected at higher external resistance (MFC-1kΩ and MFC-5kΩ). Previous studies have also reported the presence of these additional metabolites in the MFC effluents, when complex/real wastewater was used with mixed culture inoculum systems [20, 50, 83]. The prevalence of acetic and butyric acid in the MFC effluents might be due to the abundance of acetic and butyric producers (*Clostridia* and *Bacteroidia*) in the enriched anodic inoculum [21, 51]. The total soluble metabolite concentration increased with the external resistance, with the highest concentration observed in MFC-5kΩ. Substrate consumption rate was higher at lower external resistance, probably due to the increased electron flux to the circuit because of increased anodic respiring activity. On the contrary, higher external resistance (>1 kΩ) resulted in the substrate partial degradation, as shown in the high metabolite concentrations. This indicates a possible metabolic shift from anode respiration to fermentation. Also, the increased metabolite production resulted in a slightly acidic environment (pH 5.76 ± 0.25) relative to the lower external resistance (pH 6.2-6.5 for MFC-0.1kΩ to MFC-1kΩ. This could affect the electrogenic microbial community that in turn affects the electron transfer process [80]. Despite MFC-0.1kΩ exhibiting high substrate conversion, the system still accumulated soluble metabolites of 2,667 ± 288 mg/L. This suggests that oxidation of the organic content of DFPOME effluents is probably the rate-limiting step in the anaerobic degradation process. Thus, optimizing the substrate concentration could potentially improve substrate utilization.

#### 3.6.2 Anolyte pH

Several different anolyte pH was investigated while the catholyte solution was maintained at pH 7. Undiluted DFPOME was the electron donor and operated with 1 kΩ external resistance. MFC-pH5 had the lowest voltage generated compared to the MFCs with anolyte adjusted to neutral and alkaline pH (Fig. 8a). The maximum voltage generated with the different anolyte adjusted to different pHs were as follows: 182.82 ± 21.56 mV (pH 5), 409.01 ± 18.37 mV (pH 7), 468.23 ± 61.56 mV (pH 9), and 303.21 ± 32.45 mV (pH 11). A similar trend was observed with the power and current densities (Fig. 8b), with MFC-pH5 exhibiting the lowest current and power density output of 69.51 ± 6.45 mA/m^2^ and 12.71 ± 3.04 mW/m^2^, respectively. These outputs with acidic anolyte resulted in about 2- and 5-fold current and power density decrease compared to operation at neutral pH. The neutral and alkaline pH (7-11) had peak power densities between 50.72-83.36 mW/m^2^ and peak current densities between 138.86-178.04 mA/m^2^. However, significant difference was observed between MFCs operated at neutral and alkaline pH (ANOVA and post hoc Tukey test; P < 0.05). MFC-pH9 showed the highest power and current density of 83.36 ± 6.18 mW/m2 and 178.04 ± 17.2 mA/m^2^, respectively, which is a 1-fold increase compared to that at neutral pH. A further increase in anolyte pH to 11 decreased the MFC performance.

**Fig. 8.**
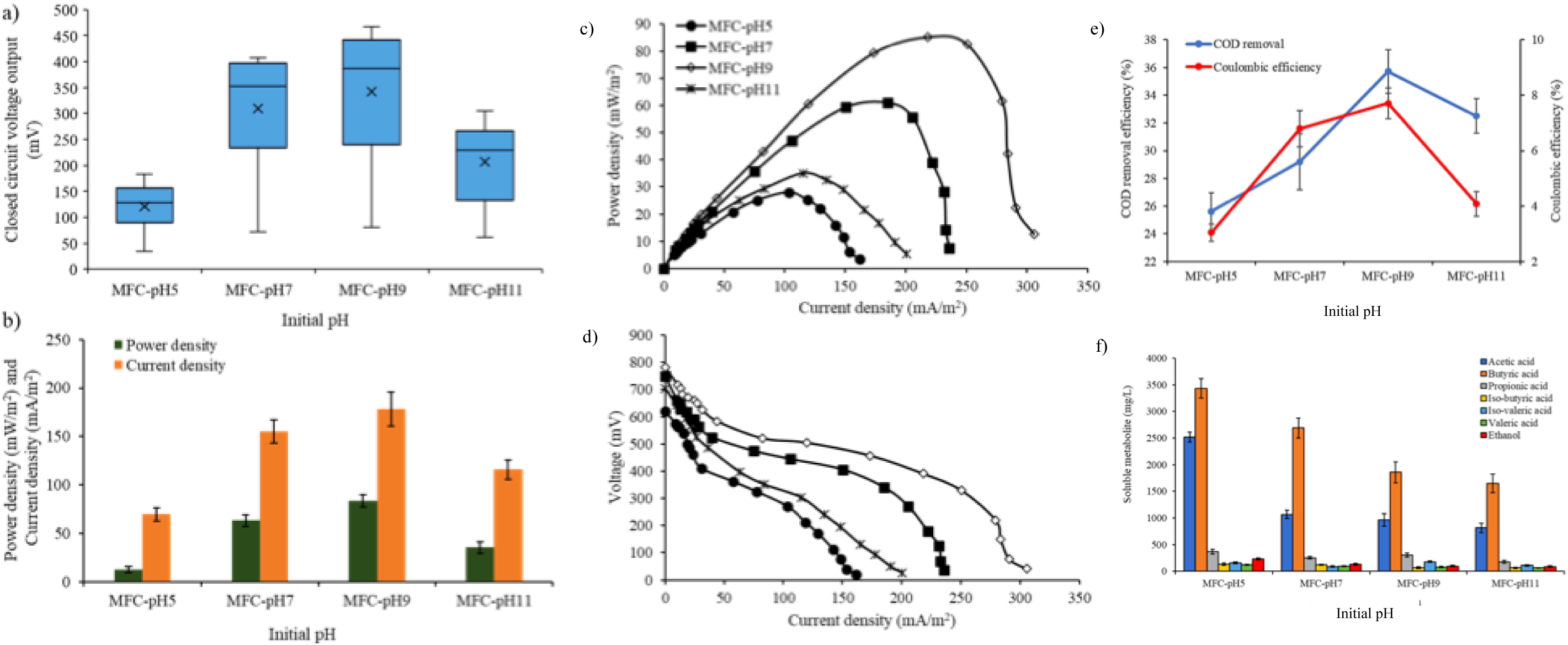
MFC power performance under the different initial anolyte pH (5, 7, 9, and 11), (a) closed circuit voltage, (b) maximum power and current density, (c) power density-current curves, (d) voltage-current curves, (e) organic matter (COD) removal and Coulombic efficiencies, and (f) metabolite profile

Anolyte pH is an important parameter in the MFC system, as it influences the bacterial metabolisms, generation of electrons and protons, and the kinetics of proton transport from the biofilm to the cathode chamber [84]. As such, pH is normally regulated using a buffered solution, or acidic/alkaline solution for the buffer-less system, to facilitate the respective reactions. In this current study, a buffer-less anolyte with the pH adjustment was carried out with the addition of dilute acid (1 M HCl) and alkaline (1 M NaOH) solutions. The generation and accumulation of protons result in the acidification of the anode chamber environment. This is because the generation and consumption of protons occur simultaneously, and the transfer of protons via the membrane may not be at the same rate as the electron transfer via the circuit. Thus, the anolyte pH drop is inevitable, especially with organic-based substrates. In addition to the two reactions (electron generation and proton transfer), other metabolic reactions (e.g. metabolite production) also occur, decreasing the pH of the anodic chamber [44]. Similar to previous studies [47, 85], this study shows that neutral and alkaline conditions (pH 9) were preferable for the MFC operations. pH of the anolyte decreased to 6.26 ± 0.19 (MFC-pH7) and 6.82 ± 0.21 (MFC-pH9) as the current produced. The buffering capacity of the alkaline solution and proton consumption at the cathode countered the acidification process and increased the proton flux from the biofilm and across the membrane. The increased power density at pH 9 can also be attributed to the less acidogenic activities, and more of the electrons released channeled to the current generation [45]. A previous study attributed the increased power output with alkaline environments to the anode electrodes exhibiting more negative potentials [86]. The results of this current study align well with several previous studies with MFC systems treating complex waste, reporting an optimum pH in the range of 8-10 [47, 87].

A further increase of the anolyte pH to 11 affected the electrochemical activity of the anode biofilm. The decreased proton concentration resulted in poor proton gradients across the membrane, limiting the cathodic reactions [44, 49]. This explains the 1.3-fold decrease in the power output compared to MFC-pH9. However, the power output at pH 11 was higher than that of pH 5, suggesting the better adaptation of the MFC microbial community to an alkaline environment. The decrease in power output at pH 11 contrasts with a previous study [86], which reported higher power density when the anolyte was adjusted to pH 11. However, the biofilm in the previous study was developed under a much higher anolyte pH of 10 compared to this study, allowing a more alkaline-adaptable community. Overall, the MFCs operated at pH 7 and 9 provide a suitable condition for the growth and activity of electrogens [88, 89]. The peak power densities measured from the polarization analysis also showed better performance as the pH of the anolyte increased to pH 9 (Fig. 8c). Further increase to pH 11 did not show improved power density, as described above. The internal resistance calculated from the voltage curves (Fig. 8d) also showed that adjusting the pH to 9 decreased the internal resistance of the system, with MFC-pH9 having the lowest internal resistance of 434 Ω. MFC-pH5 and MFC-pH11 had high internal resistances of 909 Ω and 800 Ω, respectively. The low internal resistance with increased anolyte pH aligns with the previous report [47] regarding low activation loss, as the alkaline anodic environment may have facilitated the establishment of an electrogenic bacterial community. Higher internal resistance with acidic anolyte of pH 6.0 and 5.0 respectively, has also been reported, compared to the operations at higher pH values [89, 90].

The highest COD removal efficiency of 35 ± 2.61% was observed with MFC-pH9, and the removal efficiency decreased as the anolyte pH shifted towards acidic and more alkaline conditions (Fig. 8e). MFC-pH5 had the lowest COD removal efficiency of 25.61 ± 1.87%, about 1.4- and 1.1-fold decrease in removal rate compared to that of MFC-pH9 and MFC-pH7, respectively. Under acidic condition, metabolism is typically directed towards metabolite production, and possibly hydrogen production, decreasing the organic matter removal rate [45]. Higher COD removal at pH 7 and 9 is accompanied by increased current production. Previous studies attributed the higher COD removal rates at neutral pH to the proliferation of methanogenic activities [91]. However, in this current study, methanogenic activities were suppressed. Thus, the increased COD removal can instead be attributed to the increased electron transfer rate, which resulted in increased substrate metabolisms. Despite the higher COD removal (32.5 ± 1.25%) with MFC-pH11 compared to MFC-pH7, the electrochemical performance was lower. This implies that COD utilization was more for cellular growth and maintenance rather than for the current generation. Likewise, the Coulombic efficiency (CE) increased from 3.04 ± 0.32% to 7.71 ± 0.56% when the anolyte pH increased from 5 to 9. Further increment to pH 11 decreased the CE by 1.9-fold, suggesting less extracellular electron transfer at acidic and strong alkaline environments.

Metabolite profiling showed that neutral and alkaline conditions have lower metabolite concentrations than the acidic condition, suggesting that the acidogenic bacteria were more active in pH 5 (Fig. 8f) [92]. The results correlate with the COD removal results, in which higher COD removal and lower metabolite concentrations were observed with neutral and alkaline anolyte pH, while MFC-pH5 had the highest metabolite concentrations of 6972.2 ± 332 mg/L. The decrease in metabolite concentration as the anolyte pH increase suggests the decrease in acidogenic activities at these conditions, as reflected in the resulting pH of the effluents 6.26 ± 0.19 (MFC-pH7) and 6.82–7.85 (MFC-pH9 and MFC-pH11). Thus, anaerobic respiration was more prominent and resulted in increased MFC performance. Under the acidogenic condition, complete substrate oxidation is not feasible, and most of the electrons are channeled to metabolite production rather than for anaerobic respiration. This correlates with the decreased current production observed with the acidic condition.

#### 3.6.3 DFPOME Concentration

MFC performance was investigated at different DFPOME organic loads: 25% (4,220 mg/L COD), 50% (8,500 mg/L), 75% (12,750 mg/L), and 100% (undiluted) (17,300 mg/L). The anolyte pH was maintained at 7, with 1 kΩ external resistance used. When the DFPOME was diluted, the generated voltage was much lower with 25% DFPOME (89.90 ± 16.58 mV) and 50% DFPOME (245.07 ± 23.42 mV) compared to undiluted DFPOME (409.01 ± 18.37 mV) (Fig. 9a). In contrast, operations with 75% DFPOME concentration generated a maximum voltage output of 457.44 ± 33.55 mV, significantly higher (T-test, P < 0.05) than with undiluted DFPOME. A shorter cycle duration of four and six days was also observed with MFC-25% and MFC-50% operations compared to the longer cycle duration of nine days with MFC-75% and MFC-100%. This suggests faster degradation and utilization of the DFPOME with lower initial COD concentrations. MFC-75% has the highest power density of 79.56 ± 5.31 mW/m^2^ and current densities of 173.93 ± 12.49 mA/m^2^ (Fig. 9b), which was a 1-fold increase to the power output with MFC-100%. Lower power and current densities were observed with MFC-25% (3.07 ± 0.76 mW/m^2^, 34.18 ± 5.13 mA/m^2^) and MFC-50% (22.83 ± 7.51 mW/m^2^, 93.18 ± 10.14 mA/m^2^).

**Fig. 9.**
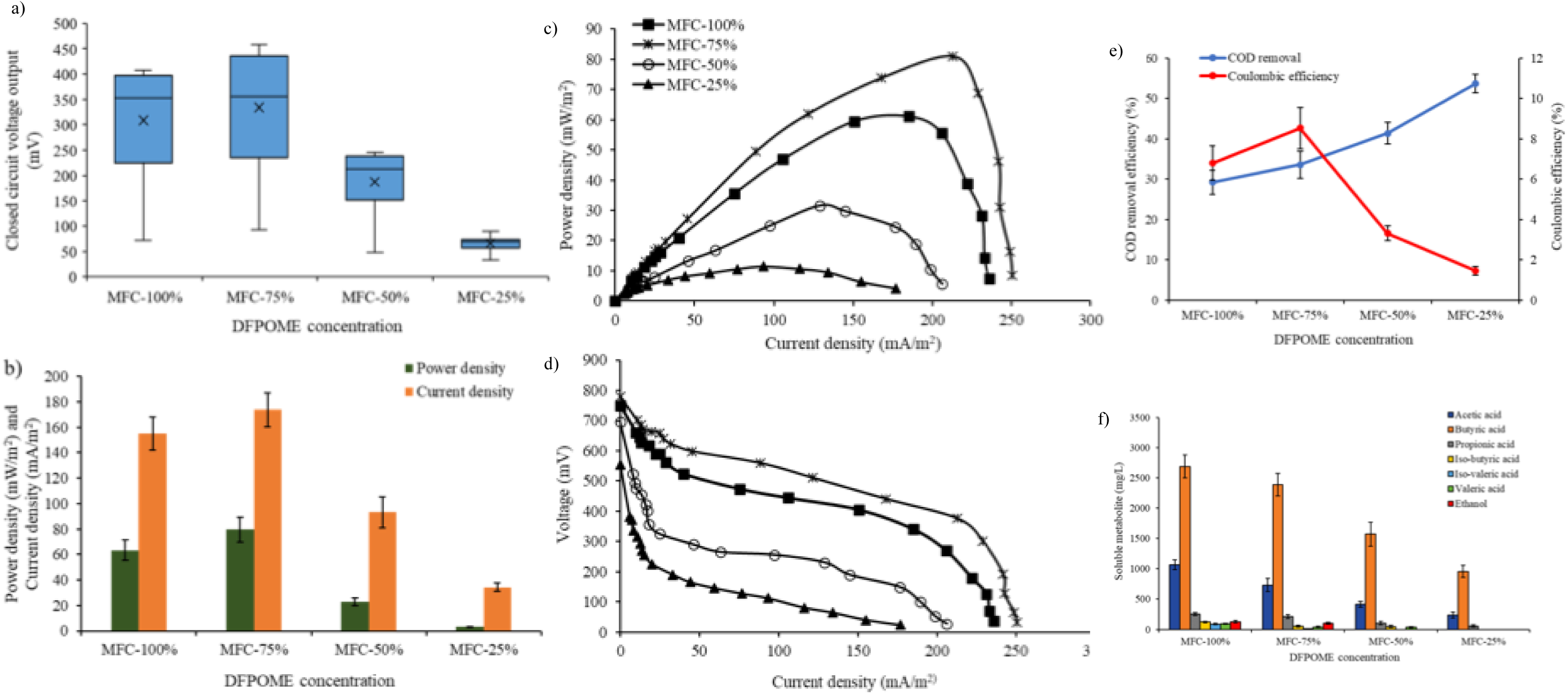
MFC power performance under the different DFPOME concentrations (100%, 75%, 50%, and 25%), (a) closed circuit voltage, (b) maximum power and current density, (c) power density-current curves, (d) voltage-current curves, (e) organic matter (COD) removal and Coulombic efficiencies, and (f) metabolite profile

Ionic strength of the substrate used as an electron donor is an important factor driving the biological functions affecting the power performance [93]. The oxidation rate depends on the concentration of the substrates [45]. High organic load in the undiluted DFPOME showed high power production. However, dilution to 75% improved this further. From then onwards, lower organic content with further dilution resulted in decreased power production, potentially due to the lower metabolic activities. A similar outcome was reported before, in which undiluted POME effluents resulted in a power density of 2-fold lower than operation with 50% POME effluent using MnO_4_ catalyzed cathode electrode and pure-culture inoculum [40]. Other studies also reported decreased power output as the organic content decreased [21, 22, 29, 47].

Polarization measurements also increased power output with substrate dilution concentration to 75% DFPOME (Fig. 9c,d). The internal resistance decreased as the substrate concentration decreased (increased dilution), with the following values: 486 Ω (MFC-25%), 434 Ω (MFC-50%), 506 Ω (MFC-75%), and 543 Ω (MFC-100%). Lower internal resistance at lower substrate concentration can be attributed to the less interrupted flow of electrons to the anode. However, more substrates were available for the electrochemical processes at higher substrate concentrations, suggesting the importance of substrate dilution for optimum MFC operation.

COD removal efficiency was higher as the substrate concentration decreased, accompanied by the decrease in current production (Fig. 9e). The maximum COD removal efficiencies for each substrate concentration were 53.69 ± 2.33% (MFC-25%), 41.42 ± 2.64% (MFC-50%), 33.58 ± 3.37% (MFC-75%), and 29.22 ± 5.01% (MFC-100%). The amount of COD oxidized at 75% substrate concentration (4852.31 ± 365.5 mg/L) was almost twice that of MFC-25% (2587.86 ± 245 mg/L), despite the lower COD removal efficiency. This suggests higher concentrations of biodegradable substrates were available for current production at 75% DFPOME concentration, as observed previously [22]. Coulombic efficiency, on the other hand, increased with the increase in DFPOME concentrations, from 1.47 ± 0.22% (MFC-25%) to 8.53 ± 1.12% (MFC-75%). This suggests that at low organic content, most of the electrons from the substrate oxidation were consumed in other microbial metabolic functions rather than for current generation. In general, 75% DFPOME concentration was found to be the optimum concentration, based on power production. The final metabolite profiles of the runs at different substrate concentrations (Fig. 9f) showed higher soluble metabolite concentration at high substrate concentrations, suggesting that higher organic load provides more substrate for conversion. A similar was reported before, where VFA concentration increased with the increase in the influent COD concentration of the synthetic wastewater used [94].

#### 3.6.3 Serial and parallel configurations

Based on the outcome of the single-factor optimizations, a set of optimum conditions was obtained from the factors which yielded the maximum power output in each parameter studied. The conditions selected were (0.5 kΩ, pH 9, and 75% DFPOME concentration) based on the improved electricity generated and higher wastewater treatment efficiency. The repeated batch operation yielded power and current densities between 79.17-93.09 mW/m^2^ and 245.37- 263.59 mA/m^2^. The relatively stable period was about 82 h, and about 1-fold increase was achieved for power and current density compared to the acclimatized condition, as summarized in Table 2. COD removal and Coulombic efficiency also increased to 36.17 ± 2.25% from 29.22 ± 5.01% and 8.53 ± 1.35% from 6.80 ± 0.66%, respectively. The optimum conditions improved the current generation and power density, higher than the previously reported output using mixed culture inoculum [22, 24, 29].

**Table 2.** Comparison of the power performance from the acclimatized and optimum conditions in this study and other MFC studies using mixed culture inoculum.

| Conditions | Power Density<br>(mW/m<sup>2</sup>) | Current Density<br>(mA/m<sup>2</sup>) | COD Removal<br>(%) | Coulombic<br>Efficiency (%) | Reference |
| --- | --- | --- | --- | --- | --- |
| (1 kΩ, pH 7, 19,300 mg/L) | 63.31 ± 8.07 | 155.16 ± 12.88 | 29.22 ± 5.01 | 6.80 ± 0.66 | Acclimatized (This study) |
| (0.5 kΩ, pH 9, 14,450 mg/L) | 93.09 ± 11.52 | 263.59 ± 19.32 | 36.17 ± 2.25 | 8.53 ± 1.35 | Optimized (This study) |
| (1 kΩ, pH 7, 1,000 mg/L) | 8.34 (W/m<sup>3</sup>) | 263 | 64 | 23 | [21] |
| (0.5 kΩ, pH 7.85, and 884 mg/L) | 74 | NA | 93.57 | 10.65 | [29] |
| (10 kΩ, pH 7, 2,680 mg/L) | 85.11 | 91.12 | 13 | NA | [24] |
| (1 kΩ, pH 7, 1,000 mg/L) | 22 | NA | 70 | 24 | [37] |

The power output of the optimum condition in a single cell was still considered insufficient to be used to power electrical devices. Combining single cell units in parallel and serial connections could provide the required power and current density. For this purpose, the performance of three MFC units connected in series and parallel was investigated. The connected electrical configurations were initially operated in OCV mode, followed by varying external resistance (polarization). The OCV for the individual MFC units varied between 790-810 mV, while that of the stacked MFC units was 805 mV (parallel) and 2,380 mV (serial). The OCV of the serial circuit is the cumulative voltage generated from the individual MFC units (795, 800, and 780 mV).

Polarization measurement showed that the maximum power density that could be obtained was at an external resistance of 300 Ω (parallel) and 1600 Ω (serial). This generated a power density of 142.63 mW/m^2^ (425.76 mA/m^2^) and 123.35 mW/m^2^ (171.21 mA/m^2^) for parallel and serial circuit connections, respectively (Fig. 10a,b). The connected MFCs were then operated semi-continuously by replacing 75% of feed every 82 h for a period of one month.

**Fig. 10.**
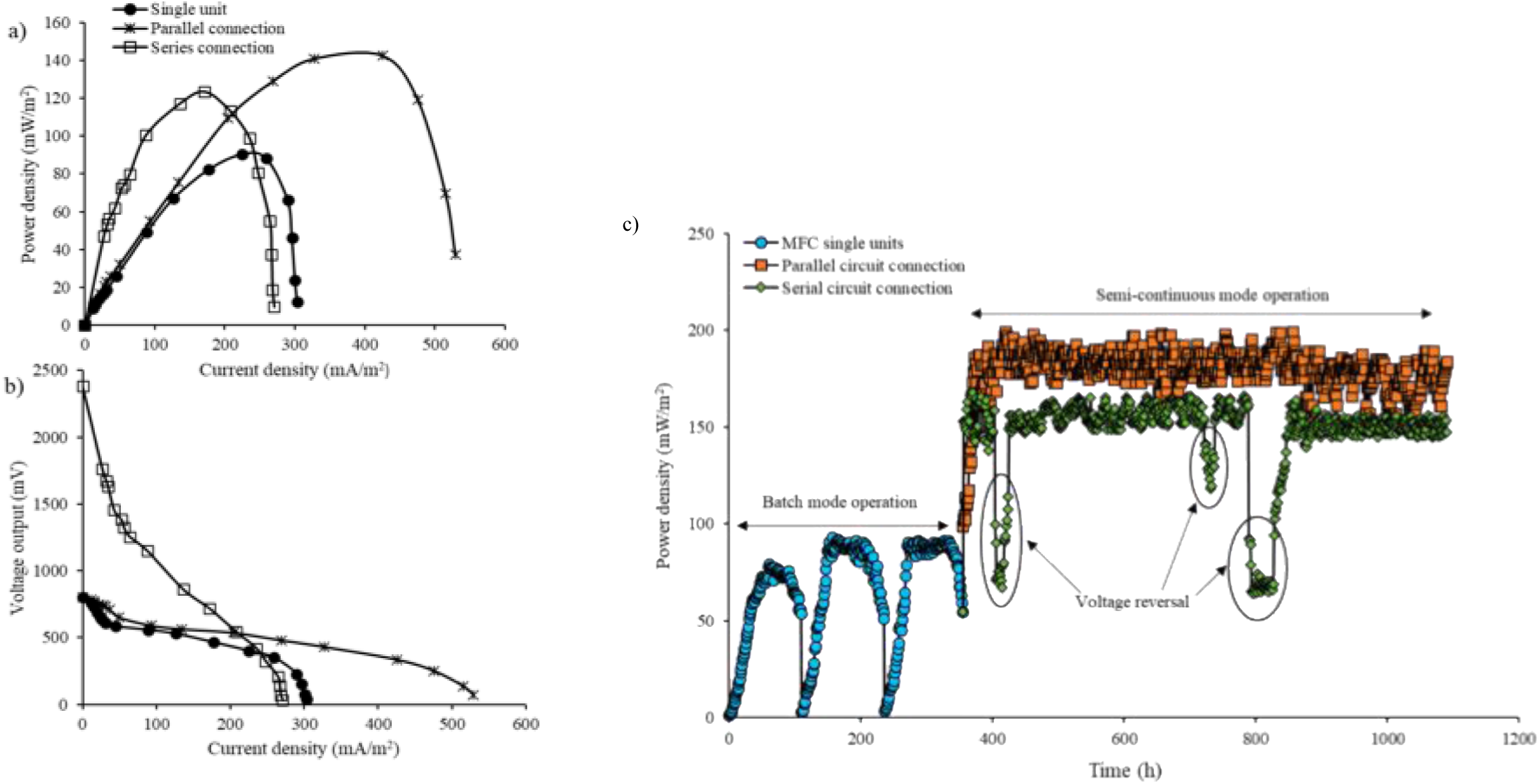
MFC power performance under different MFC configurations, (a) power density-current curves, (b) voltage-current curves, and (c) power density output during a 30-day semi-continuous mode of operation

The semi-continuous mode of operation with the stacked MFC configurations produced a relatively stable power production with the parallel circuit connection, as shown in Figure 10c. The average power and current densities obtained were 174.35 ± 22.47 mW/m2 and 469.55 ± 17.06 mA/m^2^. The power and current output was almost twice higher than that of the single units. Power production with the serial circuit connection was not as stable, as voltage reversal occurred on days 17, 30, and 33-35, affecting the power output. The voltage reversal occurred from MFC1 (the first MFC unit in the arrangement), which exhibited negative voltage readings (−153 mV to -264 mV). This is a common occurrence with serial circuit connections, due to imbalanced potentials relating to the reaction kinetics on the anode or cathode electrode [95] and has also been reported in previous studies [96, 97]. Nonetheless, the average power density was still higher than that obtained from the single units by about a 1.5-fold increase (141.4 ± 24.12 mW/m2). This is still significantly lower (T-test: P < 0.05) than the parallel connection. The average current density (182.49 ± 17.36 mA/m2) generated was lower than the current output of the single units. The internal resistance of the parallel circuit connection was lower (330 Ω) than the serial circuit connection (1,956 Ω). This higher internal resistance probably led to the lower power output.

Successful use of MFC systems in up-scaled applications relies on implementing the system in a continuous mode of operation. Currently, there is still limited study conducted on MFC operated in a continuous mode with POME as the substrate. Most studies investigated the abilities of pure culture and mixed culture inoculum, electrode material, and different POME concentrations on power production. These studies investigated MFC operation under batch mode [22, 23, 25, 40] and fed-batch mode [21, 27]. Under these modes, the metabolic activity of the microbial community decreases as the concentration of oxidable substrates depletes after a certain period of operation.

On the other hand, several studies reported a stable operation under continuous mode. A power density of 19.79 mW/m^2^ was reported from dark fermentation effluents using a stacked MFC system with a dual gas diffusion cathode design [58]. In another study, double-chambered MFC with biocathodes was successfully operated continuously for 6 months for the treatment of swine manure, achieving a power density of 2-4 W/m^3^ [97]. A 12-month continuous operation using an aerated buffered catholyte was conducted with raw POME diluted with a phosphate-buffered solution [26]. The substrates utilized in these studies all had lower COD concentrations (less than 3,000 mg/L). This current study has shown that a relatively stable current production could be achieved with higher anolyte COD concentration. COD removal efficiency of the stacked connection was between 28.15-35.5% for the parallel connection and 25.58-31.0% for the serial connection. The higher current generated with the parallel connection could be attributed to the higher substrate conversion, as reflected in the COD removal. Higher current generation often suggests higher oxidation of the organic matter, and higher electron production due to the increased metabolic activities [98]. Stacked MFC operation showed slightly lower COD removal than the single MFC runs, probably due to the shortened retention time, decreasing the contact time between the substrate and biomass. The overall Coulombic efficiency for the stacked operations was 10.15% for parallel and 8.92% for serial circuits. The power performance of the stacked MFC connections is summarized in Table 3. Our results indicate that the performance of a single MFC unit improved when in stacked arrangements, in agreement with previous studies [97, 99]. Although the serial circuit connection had an issue with voltage reversal, a mixed (series-parallel) circuit connection could potentially rectify this. Both systems combined (dark fermentation and MFC) showed a total COD removal of 72.6% for parallel circuit connection and 70.6% for serial circuit connection, higher than the individual systems.

**Table 3.** Summary of the power performance of the stacked MFC operations.

| Performance Indicator | Parallel | Serial |
| --- | --- | --- |
| Open circuit voltage (mV) | 805 | 2380 |
| Maximum power density (mW/m<sup>2</sup>) | 199.12 | 165.27 |
| Maximum current density (mA/m<sup>2</sup>) | 502.59 | 195.88 |
| Internal resistance ($\Omega$) | 330 | 1956 |
| COD removal efficiency (%) | 35.5 | 31.0 |
| Coulombic efficiency (%) | 10.15 | 8.92 |

## 4 Discussion

The integration of MFC with dark fementation offers a promising route for enhancing overall resource recovery from POME. While dark fermentation efficiently converts readily biodegradable fractions into biohydrogen, the resulting effluent remains rich in soluble organic matter, particularly volatile fatty acids, representing untapped chemical energy. Employing MFCs as a downstream polishing and energy-recovery system enables partial conversion of these residual organics into electricity, while simultaneously improving effluent quality, thereby advancing a more circular and energy-efficient POME biorefinery. However, POME-derived DF effluent is still a chemically complex matrix containing high COD, suspended solids, recalcitrant compounds, and fluctuating pH, all of which can strongly influence microbial community structure, electron transfer efficiency, and electrochemical stability. As such, systematic optimization of operational parameters is essential to balance electrogenic activity with competing metabolic pathways, and to maximize both electrical recovery and wastewater treatment performance.

During start-up operation, a clear inoculum effect was observed in our MFC system, both on the time to reach reproducible cycles and on the magnitude of power generation. The sludge-inoculated system (MFC-S) moved from an initially weak first cycle to progressively higher, more repeatable outputs, reaching reproducible performance in the later cycles with an average power density around 60 mW/m² and a maximum polarization-derived power density of 61.18 ± 6.49 mW/m². This pattern is consistent with many reports of MFC start-up, with early operation often dominated by growth and attachment of electroactive consortia, after which electron transfer becomes more efficient as anode biofilms mature and internal losses stabilize [100].

By contrast, the system without added sludge (MFC-WS) produced much lower and less stable power, with sharper declines and shorter apparent “stable” periods, despite achieving COD removal in a similar range. This divergence between COD removal and electrical recovery is a common observation for complex wastewaters and mixed communities: a large fraction of electron equivalents can be diverted to fermentation products, biomass synthesis, or competing respiratory pathways rather than being captured as current. This is supported by the more acidic effluent pH for MFC-WS, indicating accumulation of volatile fatty acids and non-electrogenic metabolism. It is known that wastewater COD removal does not directly translate to Coulombic efficiency (CE), especially under mixed-culture, non-ideal electron-acceptor conditions [100].

During acclimatization, MFC-S achieved a maximum COD removal of 29.22 ± 2.01% with a maximum CE of 6.89 ± 0.89%, while MFC-WS reached 22.98 ± 2.84% COD removal and a lower maximum CE (about 1.5%). These outputs are anticipated for acidogenic, fermentation-derived effluents, where VFAs and soluble intermediates are abundant but microbial competition remains strong. The low CE is particularly expected when multiple sinks exist for reducing equivalents, and when the substrate matrix contains components that are not readily accessible to anode respiration without extensive hydrolysis or syntrophic conversion. The performance profile supports the potential integration of dark fermentation with downstream bioelectrochemical systems. Acidogenic effluents are often VFA-rich and therefore suitable for MFC feeding. MFC-S displayed a more interpretable linear region and achieved higher current densities than MFC-WS, which showed an early sharp voltage drop consistent with higher activation losses or weaker catalytic kinetics [100].

The running parameter optimization indicates that initial pH adjustment to alkaline conditions (pH 9) increased performance relative to neutral pH operation, suggesting that biofilm electron transfer kinetics and charge-transfer resistance are pH-sensitive. Previously, a direct mechanistic study on anodic biofilm responses to pH found maximum power at pH 9 and attributed improvements to faster electron transfer kinetics and improved biofilm formation under alkaline conditions [101].

At the same time, alkaline conditions in real mixed communities can shift competition away from strictly acidogenic fermentation and can reduce the extent of VFA accumulation that depresses pH and inhibits electrogenesis. External resistance of 0.5 kΩ was found to be the best-performing load. A long-term study that implemented real-time optimization of external resistance demonstrated increased power output and improved CE when operating around the optimal resistance [102].

In general, external load interacts with pH and substrate strength in shaping CE, current, and COD removal [91]. A substrate concentration of 75% DFPOME produced the best overall balance of power/current generation, COD removal, and CE. This is consistent with the non-linear substrate concentration effect seen in MFCs: too low a concentration promotes fuel limitation and voltage reversal risk in stacks, while too high a concentration can increase mass-transfer constraints, elevate competing metabolism, and worsen Coulombic losses. Substrate strength has been reported to shift the balance between COD removal and electrical recovery [91].

Higher cumulative voltage/power/current output was observed for stacked configurations compared with single units, supporting that stacking addresses the voltage ceiling of individual cells. Serial stacking increases voltage, while parallel stacking increases current delivery, resulting in higher maximum power density and current density in parallel. It must be acknowledged that serial configurations are susceptible to voltage reversal when one unit becomes fuel-limited or underperforms relative to others, which can affect the net output and the biofilm formation. This has been demonstrated before and remains one of the main practical barriers to reliable series operation under variable wastewater feeding [103]. Thus, it can be established that for DFPOME, where substrate quality and biofilm activity can vary between units, parallel stacking may provide more robust power delivery, keeping in mind the voltage-reversal mitigation strategies. Overall, results from this study supported that using DFPOME as an electron donor for MFC improves effluent quality, while recovering energy as electricity. Despite the modest COD removal during acclimatization, subsequent parameter optimization and stacking configurations improve the overall performance, further supporting the feasibility.

## Conclusion

Our investigation shows that MFCs inoculated with anaerobic sludge had higher power density than MFCs without sludge inoculation. FESEM images showed the successful formation of biofilm on the anode electrodes, with an enrichment of members of *Bacillota*, *Bacteroidota*, and *Pseudomonadota*. Due to the complexity of the substrates used (POME), a synergy of metabolic cooperation of fermentative and electrogenic communities occurred to metabolize the complex substrates effectively. Although the known electrogenic bacteria were not detected in this current study, the results suggest that other electrogenic microbial communities may be involved in power generation. The optimum conditions (0.5 kΩ, pH 9, and 75% DFPOME concentration) achieved increased current production, wastewater treatment, and Coulombic efficiency compared to the non-optimized conditions. The mode of electron transport to the anode electrode was probably mainly via cytochromes mediators. Higher cumulative voltage, power, and current densities were achieved with the stacked MFC connections, compared to single MFC units. While parallel circuit connection produced higher power and current density than serial connection. This study demonstrated the feasibility of a sequential dark fermentation and MFC operation for improved effluent quality and the production of bioenergy (biohydrogen and bioelectricity) using POME as substrate.

## Declarations

The authors declared no competing interests or conflicts.

## Acknowledgments

This work was funded by Universiti Teknologi Malaysia Tier 1 Research University Grant No. 19H14, Transdisciplinary Research Grant No. 05G24, and the Tertiary Education Trust Fund (TETFUND) of the Federal Government of Nigeria.

## Authors’ contributions

Jemilatu Omuwa Audu – Conceptualization, Methodology, Data curation, Formal analysis, Software, Writing - original draft

Ng Hui Jing – Methodology, Data curation, Formal analysis, Software, Writing - original draft

Zaharah Ibrahim – Supervision, Writing – review & editing

Norahim Ibrahim – Supervision, Writing – review & editing

Wan Rosmiza Zana Wan Dagang – Supervision, Writing – review & editing

Mohd Hafiz Dzarfan Othman – Methodology

Mohd Firdaus Abdul-Wahab – Project administration, Supervision, Conceptualization, Funding acquisition

