## Supplementary Figure S1 for "Bioelectricity Generation from Acidogenic Palm Oil Mill Effluents using Microbial Fuel Cells": Supplementary Figure S1.docx

Current density output at different external resistance

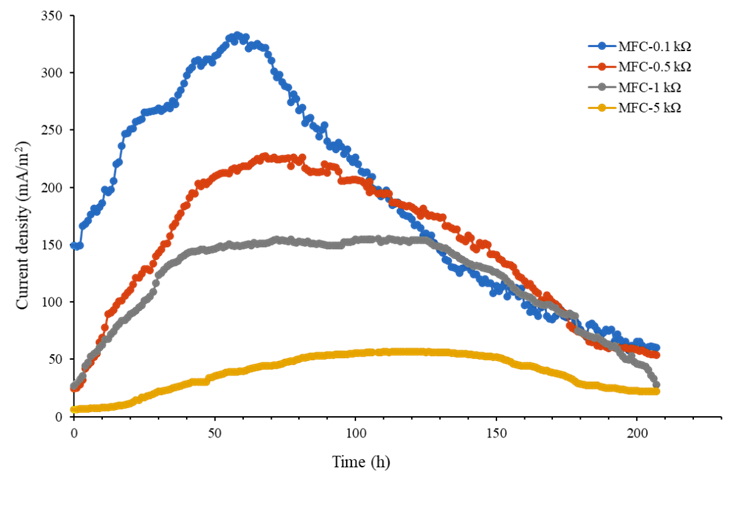

**Fig. S1** Current density output of the MFCs operated with different external resistance, showing longer stable duration with higher external resistance (1-5 kΩ)
